# Fitness Level, but not Sex, affects Exercise-Induced Pain Modulation

**DOI:** 10.1101/2025.09.16.676491

**Authors:** Janne I. Nold, Tahmine Fadai, Zora P. Gerbers, Christian Büchel

**Author notes:** Correspondence: Janne Nold, Department of Systems Neuroscience, University Medical Centre Hamburg-Eppendorf, 20246 Hamburg, Germany.

## Abstract

In a previous study by our group (Nold et al., 2025), we investigated exercise-induced pain modulation after high-intensity compared to low-intensity exercise in a heterogeneous sample of diverse fitness levels. Exploratory analyses suggested an interaction of sex, fitness level, and drug treatment, indicating that males showed increasing hypoalgesia after high- compared to low-intensity exercise with increasing fitness levels, which was diminished when naloxone was administered. In contrast, these effects were not evident in females. These exploratory findings warranted further investigation to determine if and to what extent exercise-induced pain modulation depends on fitness level and/or sex. In this current study, we investigated an all-female sample (*N* = 21) of high fitness levels using a similar paradigm as in the previous study, comparing heat and pressure pain ratings after high-intensity and low-intensity exercise. Our data show an interaction of exercise intensity and stimulus intensity in heat pain, with greater pain relief following high-intensity exercise, especially at the highest stimulus intensity. Despite results for pressure pain not reaching significance, a similar trend was evident. These results suggest that females at a high fitness level also show exercise-induced pain modulation for high-intensity compared to low-intensity exercise. Furthermore, we pooled the data of the current and previous study, providing evidence for fitness level but not sex to moderate exercise-induced pain modulation in heat pain. These findings suggest that exercise-induced pain modulation depends on fitness level but not on sex.

## Introduction

In a previous study (Nold et al., 2025), we investigated the effect of high-intensity (HI) and low-intensity (LI) aerobic exercise on pain perception in a heterogeneous sample of both sexes and diverse fitness levels (*N* = 39, 21 females), where exploratory analyses revealed a three-way interaction of sex, fitness level, and drug treatment. Males showed greater hypoalgesia after HI compared to LI exercise with increasing fitness levels. This effect was reduced when the µ-opioid antagonist naloxone was administered. These effects were not evident in females. Taken together, these results point to the nuanced interplay of individual factors such as fitness level and sex underlying exercise-induced hypoalgesia.

Repeated aerobic exercise has been shown to be associated with the activation of endogenous neuromodulatory systems implicated in pain modulation (for review see Gajic et al., 2026), such as the endogenous opioid system (Koltyn et al., 2014; Saanijoki et al., 2022) and endocannabinoid system (Matei et al., 2023). Correspondingly, previous research in humans (Geva and Defrin, 2013; Schmitt et al., 2020; Sluka et al., 2018) suggests that fitness level plays a crucial role in exercise-induced pain modulation, where subjects with higher fitness levels showed increased hypoalgesia following exercise compared to subjects with lower fitness levels (Schmitt et al., 2020). Correspondingly, higher self-reported fitness levels were shown to be associated with reduced pain ratings (Geva and Defrin, 2013), reduced temporal summation (Naugle and Riley, 2014), and greater opioid release in the orbitofrontal cortex and insula (Saanijoki et al., 2022). Crucially, most of these studies (Saanijoki et al., 2022; Schmitt et al., 2020) have been conducted in healthy all-male samples, limiting the translation to females, where the association of physical fitness and pain perception after exercise remains less clear (de Bruijn et al., 2011).

Despite the majority of chronic pain patients being female (Fillingim et al., 2009; Mogil, 2012; Rice et al., 2019), evidence on potential sex differences in exercise-induced pain modulation is surprisingly limited (Koltyn et al., 2014). Of those studies that have considered potential sex differences in exercise-induced pain modulation, some (Brellenthin et al., 2017; Koltyn et al., 2014; Niwa et al., 2022; Smith, 2004) could show that there are no sex differences, whereas others (Koltyn, 2000; Sternberg et al., 2001; Vaegter et al., 2014) suggest that hypoalgesia following exercise is more pronounced in females, potentially due to sex differences in the descending antinociceptive circuits (Fullerton et al., 2018; Mogil, 2012), but overall results remain inconclusive (Rice et al., 2019).

This current behavioural study aims to build on our previous findings (Nold et al., 2025) by investigating exercise-induced pain modulation in an independent all-female sample (*N* = 21) with a high fitness level using the same paradigm as in the previous study, without a pharmacological intervention. Importantly, fitness levels (quantified by the weight-corrected functional threshold power (FTP)) in the previous study (Nold et al., 2025) differed significantly (*P =* 0.003) between females (*M* = 1.58, *SD* = 0.44) and males (*M* = 2.03, *SD* = 0.40), thus, warranting further investigation into whether exercise-induced hypoalgesia after HI compared to LI aerobic exercise can also be observed in females with a higher fitness. Disentangling if and to what extent exercise-induced hypoalgesia depends on fitness level and sex is a crucial step to determine exercise-based interventions for (chronic) pain populations.

## Results

The final sample included *N* = 21 healthy female participants with higher fitness (weight-corrected FTP: *M* = 2.25 W/kg, *SD* = 0.41; Training volume: *M* = 13.10, *SD* = 12.28). The distribution of menstrual cycle phases (based on self-report and divided into three phases: follicular, ovulatory, and luteal as well as hormonal contraceptives) was equal across (χ2(3) = 1.66, *P =* 0.65) as well as within days (Day 1: χ2(3) = 2.26, *P =* 0.52; Day 2: χ2(3) = 0.40, *P =* 0.94; Supplemental Fig. S1A). Furthermore, the expectation of how acute exercise affects different pain domains was assessed (Lindheimer et al., 2020) and yielded no significant effects overall (joint pain: *t*(20) = -1.78, *P =* 0.09, muscle pain: *t*(20) = 0.15, *P =* 0.88). However, participants indicated that they expect acute aerobic exercise to reduce whole-body pain (*t*(20) = -2.83, *P =* 0.01; Supplemental Fig. S1B). Finally, mood ratings were assessed using the Profile of Mood States (POMS) questionnaire (Curran et al., 1995) before the experiment commenced (pre) and after it was completed (post). There were no significant differences in pre- and post-levels of fatigue (*t*(20) = -0.50, *P =* 0.62), discontent (*t*(20) = 1.84, *P =* 0.08), and drive (*t*(20) = 0.58, *P =* 0.57). However, participants rated significantly reduced dejection post compared to pre experimental testing (*t*(20) = 3.61, *P =* 0.002; Supplemental Fig. S1C).

### Successful pain calibration and exercise intervention

We calibrated pain intensity levels corresponding to 30, 50, and 70 on a Visual Analogue Scale (VAS), where VAS 0 marks the pain threshold (Supplemental Fig. S2A–B). On the experimental day, we introduced an online rating scale (real-time ratings while the stimulus is ongoing) where the pain threshold now corresponded to VAS 50 (Supplemental Fig. S2C–D) to more accurately track the temporal dynamics of pain perception through continuous ratings and incorporate non-painful sensations. Participants perceived the stimulus intensities as significantly different for pressure (*β* = 1.13, *SE* = 0.07, *t*(553.13) = 16.84, *P* < 2×10^-16^; Supplemental Table S7) and heat pain (*β* = 1.75, *SE* = 0.08, *t*(532.61) = 21.46, *P* < 2×10^-16^; Supplemental Table S9). The anticipated intensities of VAS 50 and VAS 70 were rated above the pain threshold on the online scale, indicating that they were perceived as painful (Supplemental Fig. S8), despite the ratings being lower than the anticipated intensities, potentially due to habituation effects between days (Ellerbrock et al., 2015; May et al., 2012). We achieved a significant difference between HI and LI exercise conditions in absolute power (*t*(20) = 19.34, *P =* 2.043e-14, *d* = 0.97; Fig. 1A), relative power (%FTP; *t*(20) = 42.35, *P* < 2.2e-16, *d* = 7.97; Fig. 1B), heart rate (HR; *t*(13) = 20.98, *P =* 2.073e-11, *d* = 3.45; Fig. 1C), and rating of perceived exertion (RPE; *t*(20) = 17.17, *P =* 1.942e-13, *d* = 4.61; Fig. 1D). This confirms that the exercise intensity protocols (55% FTP at LI exercise and 100% at HI exercise) elicited a different power output, relative power, physiological demand (HR), and perceived demand (RPE) and, overall, successful implementation of the exercise intervention.

**Figure 1.**
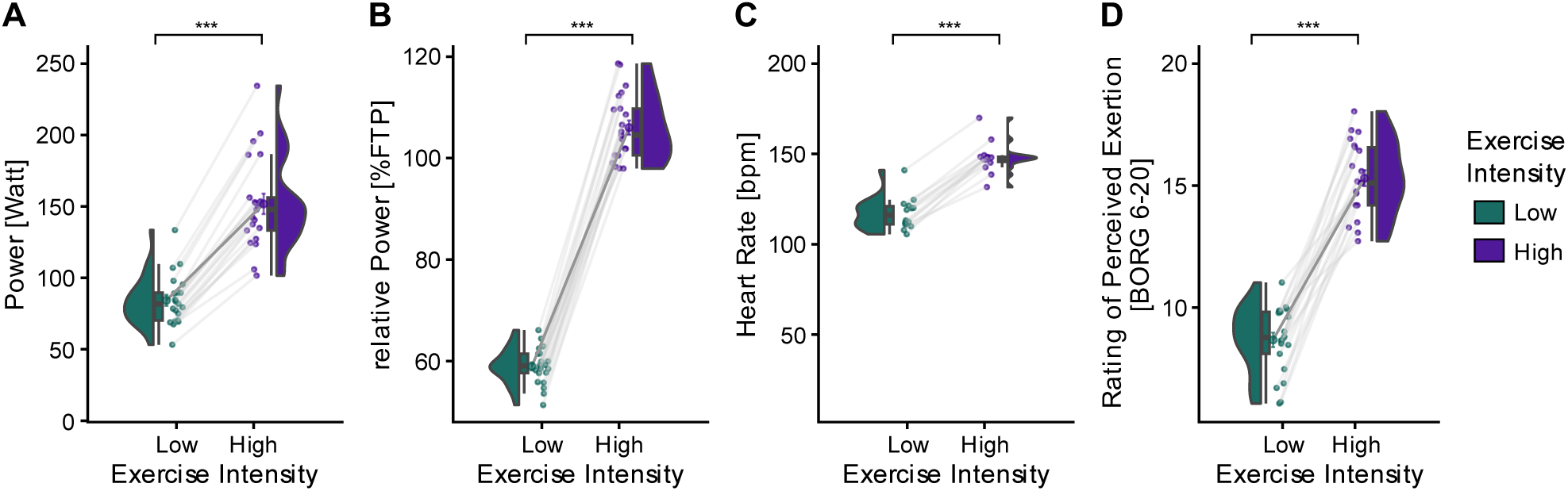
Implementation of high-intensity (HI) and low-intensity (LI) exercise. (**A**) Absolute power (Watts), (**B**) Relative power (%FTP), (**C**) heart rate in beats per minute (bpm), and (**D**) rating of perceived exertion (RPE; BORG scale) during LI (green) and HI (purple) cycling were all significantly different. *P*-values were calculated using paired *t*-tests (two-tailed, (absolute/relative) power: *N* = 21, heart rate: *N* = 14, BORG rating: *N* = 21). n.s. = not significant, * *P* < 0.05, ** *P* < 0.01, *** *P* < 0.001.

### Greater pain relief after high-intensity exercise at higher stimulus intensities

In the current study, participants provided online ratings throughout the stimulus duration. We calculated the maximum pain ratings (in the following referred to as pain ratings) across the whole stimulus duration (0 –17 seconds), which closely correspond to the pain ratings provided in the previous study (Nold et al., 2025), where participants were asked to rate the most painful sensation after the stimulus concluded. To statistically test the effect of exercise intensity (HI and LI) on pain ratings, we calculated separate Linear Mixed Effect models (LMER) with the dependent variables overall pain ratings, pressure pain ratings, and heat pain ratings, respectively. All models included exercise intensity or the interaction of exercise intensity and stimulus intensity, as well as trial and block as fixed effects, and subject as a random effect.

There was no main effect of exercise intensity on overall pain ratings (*β* = -3.29, *SE* = 2.15, *t*(1133.64) = -1.53, *P =* 0.14) as well as no main effect of exercise intensity on pressure pain ratings (*β* = -3.05, *SE* = 2.15, *t*(570.11) = -1.42, *P =* 0.16; BF_10_ = 0.46; Fig. 2A, top) or heat pain ratings (*β* = -4.13, *SE* = 2.93, *t*(538.69) = -1.41, *P =* 0.16; Fig. 2A, bottom). Bayesian model comparison showed that the model including exercise intensity was not better supported than the null model for the model across modalities (BF_10_ =0.62), as well as in pressure pain (BF_10_ = 0.64) or heat pain (BF_10_ = 0.78) only. This indicates only anecdotal evidence in favour of the null model, suggesting that the data were largely insensitive to distinguish between models with and without exercise intensity.

**Figure 2.**
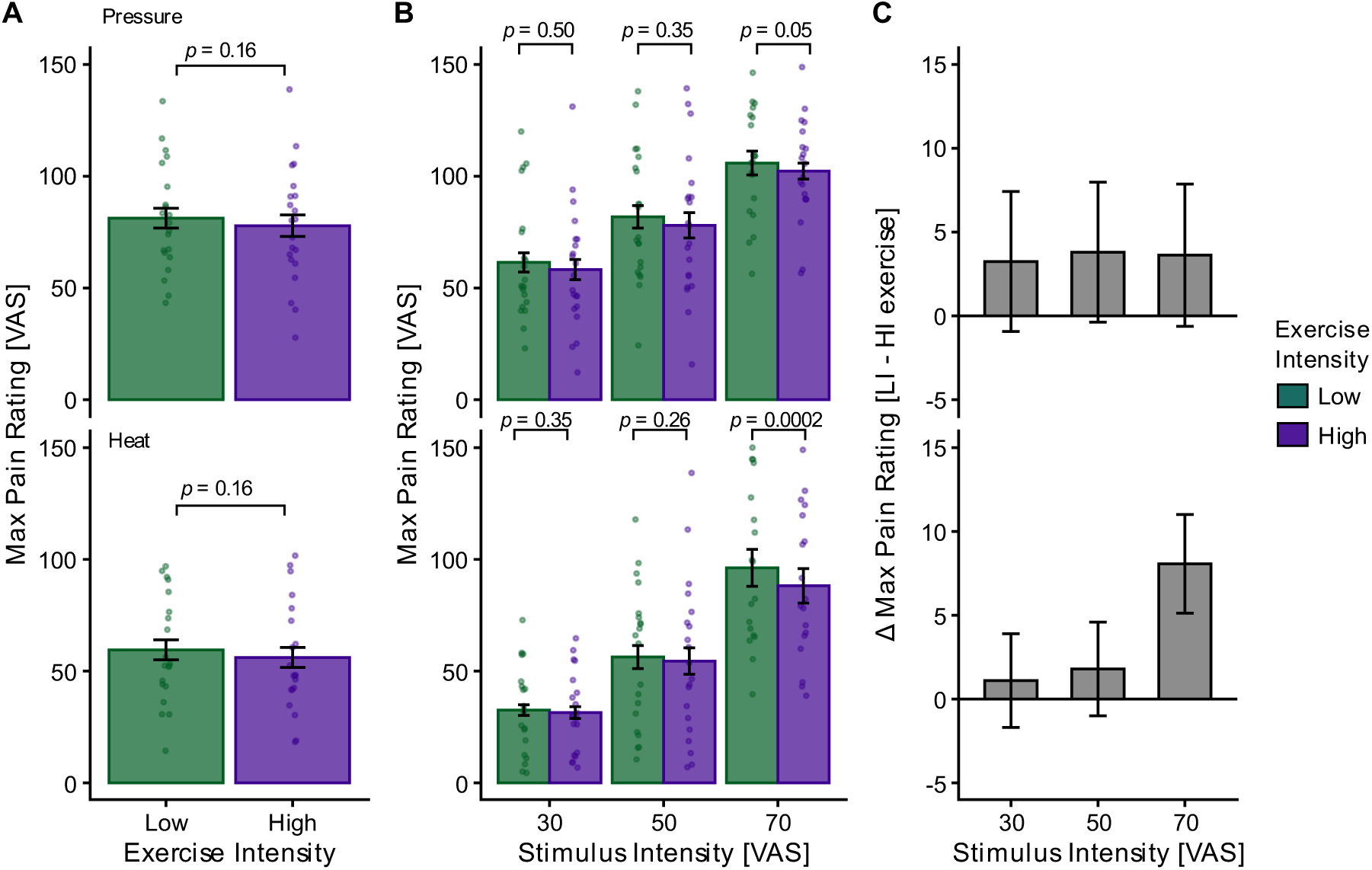
The effect of exercise intensity and stimulus intensity on pain ratings. (**A**) Pressure (top; *P =* 0.16) and heat pain ratings (bottom; *P =* 0.16) for LI (green) and HI (purple) exercise. (**B**) A significant interaction between exercise intensity and stimulus intensity was observed in heat (*P =* 0.001) but not pressure pain (*P =* 0.33). Post-hoc *t*-tests revealed that this interaction in heat pain was driven by the highest stimulus intensity (VAS 70; *P =* 0.0002). (**C**) Differences between HI and LI exercise (LI – HI) for pressure (top) and heat (bottom) pain ratings visualised at each stimulus intensity. Dots depict subject-specific pain ratings averaged across trials. *P*-values were calculated using post-hoc *t*-tests for the respective LMER models. Error bars depict the SEM (pressure: *N* = 21, heat: *N* = 20).

However, an interaction of exercise intensity and stimulus intensity on overall pain ratings was evident (*β* = -0.22, *SE* = 0.10, *t*(1119.88) = -2.22, *P =* 0.03). This interaction was driven by the interaction of exercise intensity and stimulus intensity during heat pain (*β* = -0.36, *SE* = 0.11, *t*(529.95) = -3.32, *P =* 0.001; Fig. 2B–C). Post–hoc *t*-tests revealed that the effect was especially evident at the highest stimulus intensity (*P =* 0.0002; Fig. 2B–C). For pressure pain, this interaction of exercise intensity and stimulus intensity was not significant (*β* = -0.09, *SE* = 0.09, *t*(564.99) = -0.97, *P =* 0.33; Fig. 2B–C) but a similar trend was evident when considering the post-hoc *t*-tests (Fig. 2B–C), where pain relief following HI compared to LI was greater at the highest stimulus intensity (*P =* 0.05). Bayesian model comparisons indicated evidence against an interaction between exercise intensity and stimulus intensity across modalities. For the combined dataset, the interaction model was less supported than the main-effects model (BF₁₀ = 0.22), indicating that the data were approximately 4.5 times more likely under the additive model. This pattern was consistent for both heat (BF₁₀ = 0.33) and pressure pain (BF₁₀ = 0.19), suggesting moderate evidence in favour of additive effects over interactive modulation. The full model outputs and post-hoc *t*-tests can be found in Supplemental Tables S1–S10.

### Sample differences in exercise-induced hypoalgesia

We compared the weight-corrected FTP values of the current sample with those of the previous sample (Nold et al., 2025) (*N* = 39, 21 females) for males and females separately using two-sample *t*-tests. The females employed in the current study (Fig.3A; dark blue) had a significantly higher FTP compared to the females of the previous study (light blue) (*t*(40) = - 5.00, *P =* 1.23×10^-5^; Fig. 3A). There was no significant difference in the FTP between the females in this study (dark blue) and males in the previous study (red) (*t*(36) = 1.65, *P* = 0.11; Fig. 3A). The self-reported training volume was significantly higher in the females of the current study (dark blue) compared to the females (light blue) (*t*(22) = 3.45, *P* = 0.003, Fig. 3B) and males (red) (*t*(24) = 2.82, *P* = 0.009; Fig. 3B) of the previous study. Further, analyses comparing the females of the previous study and the females of the current study concerning height, weight, and BMI revealed no significant differences but a significant age difference (*t*(32.98) = 3.27, *P* = 0.003) (Supplemental Fig. S3). Overall, this suggests that this current study included females of overall higher fitness levels compared to the previous study. In the following, females from the current study will be referred to as females with higher fitness levels due to the significantly higher FTP values than females from the previous study, as well as significantly higher training volume compared to the males and females from the previous study. These labels should be interpreted relative to the samples included in the current and previous studies rather than as absolute classifications.

**Figure 3.**
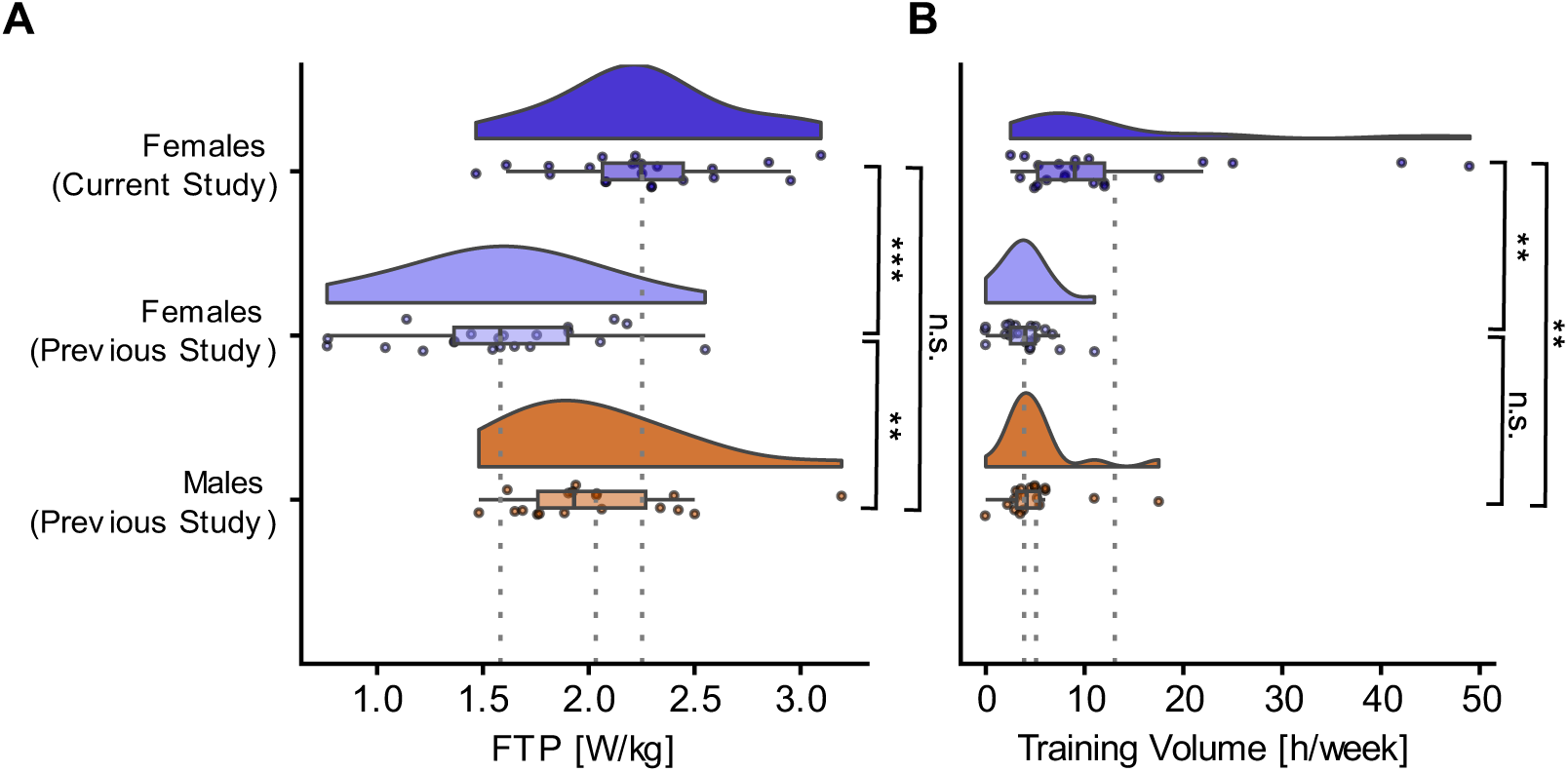
Comparing participant characteristics of the females of the current study (*N* = 21; dark blue) with the females (*N* = 21; light blue) and the males (*N* = 18; red) of the previous study (Nold et al., 2025). (**A**) Weight-corrected Functional Threshold Power (FTP Watts divided by kilograms) differed significantly between the females in the current study and the females in the previous study (*P* = 1.23×10^-5^) but not between the females in the current study and males in the previous study (*P* = 0.11). (**B**) Training volume (hours per week) differed significantly between the females in the current study and the females (*P* = 0.003) and males (*P* = 0.009) of the previous study. n.s. = not significant, * *P* < 0.05, ** *P* < 0.01, *** *P* < 0.001.

Consistent with the previous study (Nold et al., 2025), the effects in the current study were more prominent in heat pain. Therefore, all subsequent analyses for heat pain are reported in the main manuscript and for pressure pain in the supplements (Fig. S4 - S5). To allow for better comparability between the previous and current study, we rescaled the maximum pain ratings of the current study to a common scale of 0 – 100 to correspond with the previous study.

To determine whether the effect of exercise intensity differs between the participant groups contributing to the pooled dataset, we examined the interaction of exercise intensity and sub-group (males from the previous study, females from the previous study, and females from the current study). Frequentist LMERs were fitted across all stimulus intensities (VAS 30, 50, 70) and separately for VAS 70 only, as previous analyses suggested the exercise effect to be most pronounced at the highest stimulus intensity. To complement these analyses, we performed Bayesian model comparisons to quantify the relative evidence for models including exercise intensities and subgroup effects compared with simpler alternative models.

For the model across all stimulus intensities, there was no significant interaction of exercise intensity and subgroup (*β* = -1.63, *SE* = 1.68, *t*(1897) = -0.97, *P =* 0.33; Fig. 4A), with overall low absolute mean difference pain ratings between LI and HI exercise (Fig. 4B; males (previous): -0.54; females (previous): -1.74; females (current): 2.26). However, as the previous analyses revealed the effect to be most pronounced at the highest stimulus intensity, the model for VAS 70 heat pain revealed a significant interaction of exercise intensity and subgroup (*β* = -5.96, *SE* = 1.53, *t*(562) = -3.89, *P <* 0.001; Fig. 4C), where the females of the current study showed a reduced pain perception after high compare to low intensity exercise (absolute mean difference: 6.70; Fig 4D) compared to the males (absolute mean difference: -4.71; Fig 4D) and females of the previous study (absolute mean difference: -1.11; Fig 4D). The full LMER model outputs and post-hoc *t*-tests can be found in the supplements (Tables S11 – S18).

**Figure 4.**
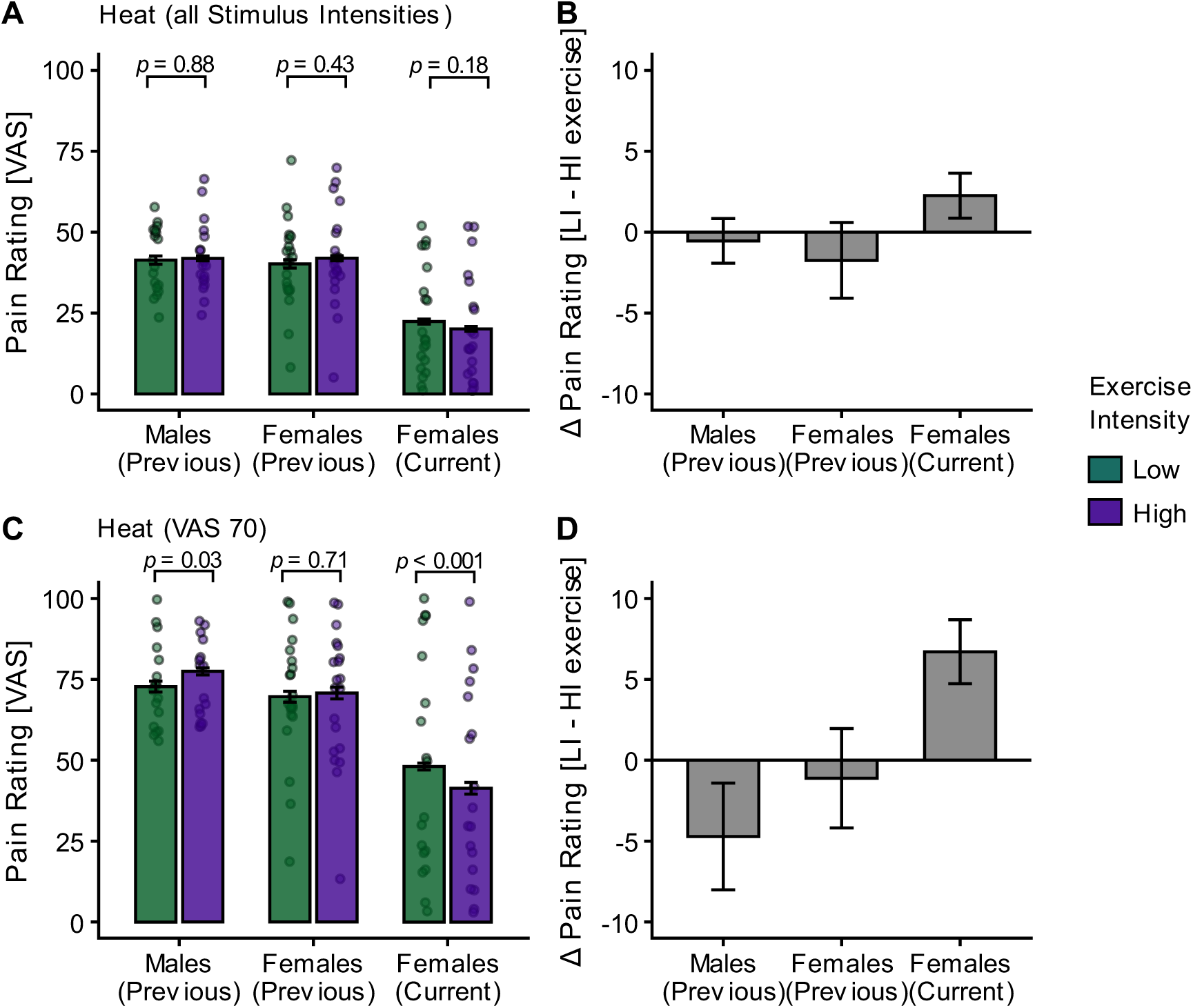
The effect of exercise intensity and subgroup (males from the previous study, females from the previous study and females from the current study) on heat pain ratings. (**A**) Heat pain ratings averaged across all stimulus intensities (VAS 30, 50, and 70) in each subgroup for LI (green) and HI (purple) exercise. There was no significant interaction of exercise intensity and subgroup on heat pain ratings (*P =* 0.33). (**B**) Differences between HI and LI exercise (LI – HI) of pain ratings across all stimulus intensities are visualised for each subgroup. (**C**) Heat pain ratings at VAS 70 in each subgroup for LI (green) and HI (purple) exercise. There was a significant interaction of exercise intensity and subgroup on heat pain ratings (*P <* 0.001). (**D**) Differences between HI and LI exercise (LI – HI) of pain ratings at VAS 70 are visualised for each subgroup. Dots depict subject-specific pain ratings averaged across trials and blocks. *P*-values were calculated using post-hoc *t*-tests for the respective LMER models. Error bars depict the SEM (*N* = 59).

Bayes model comparison for the models across all stimulus intensities indicated evidence against a model including exercise intensity alone relative to the null model (BF_10_ = 0.24). However, adding subgroup information substantially improved the model fit relative to the exercise intensity only model (BF_10_ = 1239) as well as relative to the null model (BF_10_ = 321), indicating strong evidence for an interaction between exercise intensity and subgroup. Bayes model comparison for the models including only heat pain ratings at VAS 70 mirrored these results, with evidence against the model including exercise intensity alone relative to the null model (BF_10_ = 0.22) but a substantially improved model fit when including the subgroup relative to the exercise intensity alone model (BF_10_ = 4497) and the null model (BF_10_ = 1038).

### Fitness level, but not sex, affects exercise-induced pain modulation in the pooled sample

To investigate the effects of fitness level (indicated by weight-corrected FTP) and sex on exercise-induced pain modulation, we pooled participants across studies and first conducted frequentist linear regression analyses corresponding to those used in the previous study. These analyses used participant-level difference scores (pain following low-intensity exercise minus pain following high-intensity exercise) as the outcome variable. We then performed Bayesian model comparison and Bayesian regression analyses on the pooled heat pain ratings to estimate the effects of exercise intensity, fitness, sex, and their interactions. Unlike the frequentist analyses, which focused on difference pain scores, the Bayesian models estimated posterior distributions of the regression parameters directly, yielding posterior mean estimates, 95% credible intervals (CrIs), and probabilities of effect direction without reliance on null-hypothesis significance testing (Kruschke, 2021).

The linear model including fitness level indicated a significant effect of fitness level on difference pain scores (*β* = 4.99, *SE* = 2.05, *t*(56) = 2.43, *P* = 0.02; Fig. 5A) corresponding to a correlation of *r* = 0.28, *P* = 0.03). The model including fitness level and sex also indicated a significant interaction (*β* = 11.12, *SE* = 4.69, *t*(56) = 2.37, *P* = 0.02; Fig. 5B) with a correlation of *r* = 0.57, *P* = 0.01 for males from the previous study (Fig. 5B, red line): and a non-significant correlation for females pooled across the previous study and current study (Fig. 5B, pink line) of *r* = 0.17, *P* = 0.28. The model including fitness level and subgroup also revealed a significant interaction (*β* = -7.99, *SE* = 2.76, *t*(56) = -2.89, *P* = 0.005; Fig. 5C), with weak and non-significant correlations for females from the previous (*r* = 0.08, *P* = 0.72; Fig. 5C, light blue line) and current study (*r* = -0.15, *P* = 0.54; Fig. 5C, dark blue line). The full LMER model outputs can be found in the supplements (Tables S19 – S21). Corresponding analyses using training volume as a measure for fitness level have been included in the supplements (Fig. S6).

**Figure 5.**
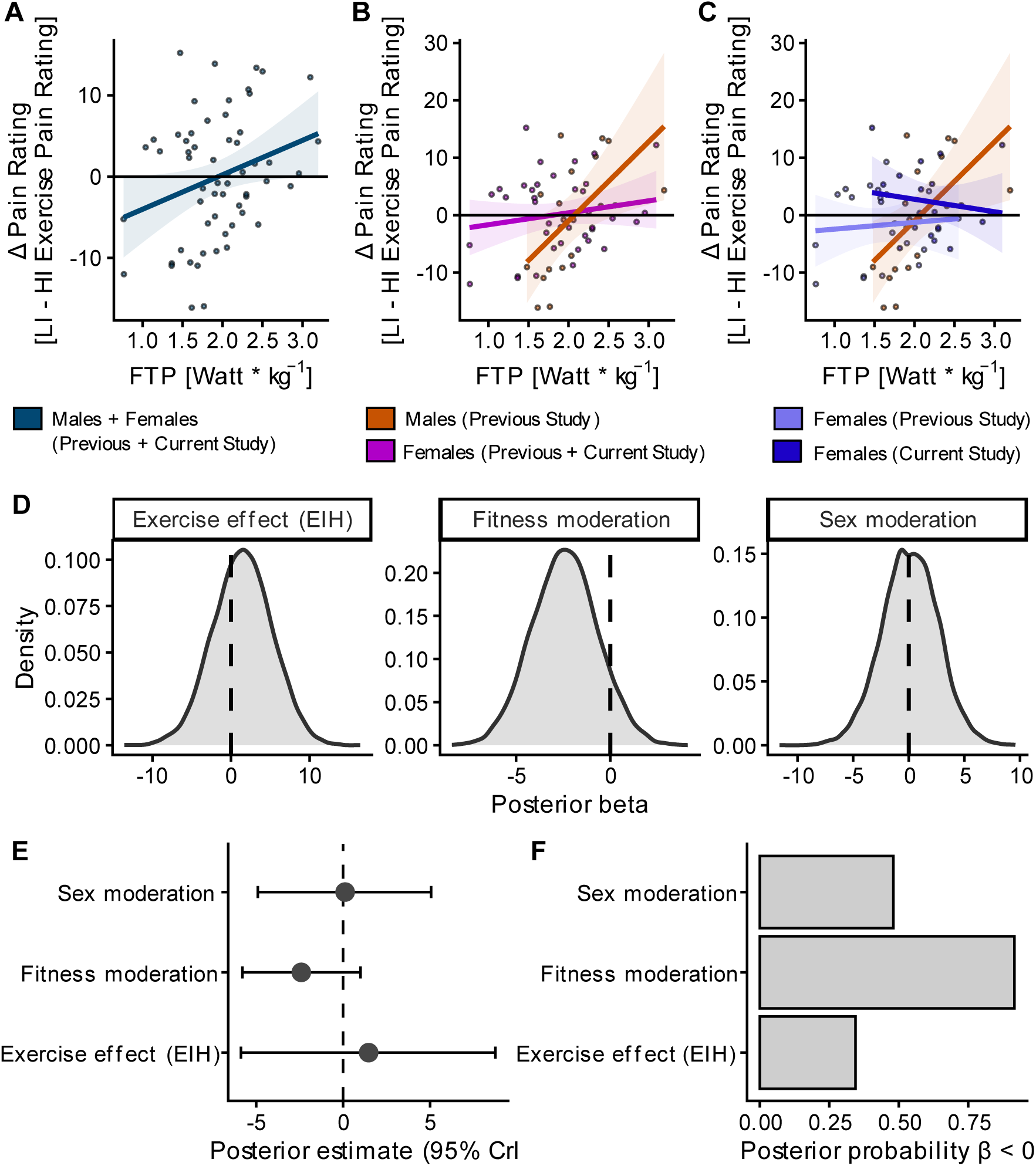
Frequentist and Bayesian analyses of exercise intensity, fitness level and sex on heat pain across pooled study samples. (**A**) Subject-specific differences in heat pain ratings (dots) between low-intensity (LI) and high-intensity (HI) exercise conditions (LI – HI exercise pain ratings) and corresponding regression line pooled across all stimulus intensities. Fitness level (measured as functional threshold power; FTP) showed a significant main effect on difference pain ratings (*P* = 0.02) and a positive relation to heat pain ratings (*r* = 0.28, *P* = 0.03). (**B**) A significant interaction of fitness level × sex on difference pain ratings (*P* = 0.02) with a significant relation in males (red line; *r* = 0.57, *P* = 0.01) but not in females (pink line; pooled across current and previous study; *r* = 0.17, *P* = 0.28). (**C**) Significant interaction fitness level × subgroup (*P* = 0.005) with non-significant correlations for females from the previous (r = 0.08, P = 0.72; light blue line) and current study (*r* = -0.15, *P* = 0.54; dark blue line). (**D**) Posterior density distribution, (**E**) posterior mean estimates and 95% Credible Intervals (CrI) and (**F**) posterior probability for the effect of exercise intensity (*β* = 1.50, 95% CrI [-5.88, 8.76], *Pr(β < 0)* = 0.34), fitness level moderation (*β* = -2.41, 95% CrI [-5.79, 1.00], *Pr(β < 0)* = 0.92) and sex moderation (*β* = 0.13, 95% CrI [-4.91, 5.06], *Pr(β < 0)* = 0.48) on heat pain ratings. The dashed vertical line indicates the null effect (*β* = 0), corresponding to no association between the predictor and pain ratings.

For the Bayesian hierarchical models, model comparison indicated evidence against a model containing exercise intensity only relative to the null model (BF_10_ = 0.24). Adding fitness (BF_10_ = 1.61) or sex (BF_10_ = 1.48) provided only anecdotal improvements over the model, including exercise intensity only. The inclusion of study (“previous”, “current”) substantially improved the model fit (BF_10_ = 2454). Finally, the full model was strongly favoured over both the exercise intensity only model (BF_10_ = 644) and the null model (BF_10_ = 158).

For the exercise intensity effect, posterior mean estimates indicated substantial uncertainty regarding the influence on heat pain ratings (*β* = 1.50, 95% CrI [-5.88, 8.76]; Fig. 5D–E) and a posterior probability of a negative effect of *Pr(β < 0)* = 0.34 (Fig. 5F). The posterior distribution for the interaction of exercise intensity × fitness level was predominantly negative (*β* = -2.41, 95% CrI [-5.79, 1.00]; Fig. 5D–E) with a posterior probability of 92% (*Pr(β < 0)* = 0.92) of a negative effect (Fig. 5F). Although the credible interval included zero, the posterior distribution suggested a probable negative moderation effect of fitness on the relationship between exercise intensity and fitness level on heat pain ratings. The tendency toward a negative exercise intensity × fitness level interaction is compatible with the frequentist analyses showing that higher fitness was associated with greater exercise-induced analgesia (Fig. 5A), although the two analyses address different model parameters and should not be interpreted as directly equivalent. Finally, the posterior estimates for the exercise intensity × sex interaction provided no meaningful evidence for an effect (*β* = 0.13, 95% CrI [-4.91, 5.06]; Fig. 5D–E), with a posterior probability of *Pr(β < 0)* = 0.48 (Fig. 5F), indicating equal support for positive and negative effects.

## Discussion

In this study, we investigated exercise-induced pain modulation between high-intensity (HI) and low-intensity (LI) exercise in a sample of females (*N* = 21) of relatively higher fitness levels. In a previous study by our group (Nold et al., 2025), exploratory analyses suggested a three-way interaction of fitness levels, sex, and drug treatment, where males showed hypoalgesia after HI compared to LI exercise with increasing fitness levels, which was diminished when naloxone was administered. These effects were not evident in females. In this current study, we aimed to explore whether exercise-induced pain modulation is also evident in females of higher fitness levels and whether this is moderated by fitness level or sex. We could show that, for heat pain, females from the current study showed pain relief after HI compared to LI exercise, especially for highly painful stimuli. A similar trend was observed for pressure pain. To place these findings in the context of our earlier observations, we pooled data from the current and previous studies and examined the influence of fitness and sex on exercise-induced pain modulation. Across the combined sample, both frequentist and Bayesian analyses indicated that fitness was associated with exercise-induced hypoalgesia, whereas there was little evidence for sex-dependent effects. Together, these findings provide support for a role of fitness level in explaining variability in exercise-induced pain modulation, while offering little evidence that exercise-induced pain modulation differs systematically between males and females.

### Effect of exercise and stimulus intensity on pain in the current sample

Corresponding to our previous study (Nold et al., 2025), there was no overall effect of exercise intensity on heat or pressure pain in the current study. However, we identified an interaction of stimulus intensity and exercise intensity that was driven by heat pain, with greater pain relief after HI compared to LI exercise at high stimulus intensity (VAS 70). A similar trend was evident in pressure pain, but it did not reach significance. These findings align with the results of our previous study (Nold et al., 2025), where exercise-induced pain modulation after HI compared to LI exercise was more prominent in heat than pressure pain.

### Pooled analyses of the current and previous study sample (Nold et al., 2025)

We confirmed that the females of the current sample showed significantly higher fitness level (as measured in weight-corrected FTP) than the females of the previous study (Nold et al., 2025). Furthermore, the absolute training volume was significantly higher compared to the males and females of the previous study, suggesting overall higher fitness levels of the females of the current study. Interestingly, although the training volume of the females of the current study was significantly higher compared to the males of the previous study, the FTP did not significantly differ, which suggests the FTP to be a more sensitive measure for fitness level compared to self-reported training volume. Previous research has shown that the FTP and FTP20 tests are reliable and convenient methods to estimate approximate measures of VO_2max_ (Denham et al., 2020) and that the FTP test is a useful test for performance prediction in moderately trained cyclists (Sørensen et al., 2019). Furthermore, as pain is a dynamic experience, we utilised real-time pain ratings in the current study as opposed to static post-stimulus pain ratings (Nold et al., 2025) to better capture the temporal dynamics of pain. Previous research (Koyama et al., 2004) has shown that, in heat pain, real-time ratings account for *R^2^* = 0.89 of the variability of post-stimulus rating, where much of the variability for the post-stimulus intensity pain rating is accounted for by the peak response during the online ratings (adj. *R^2^* = 0.3 – 0.4), suggesting good comparability of both pain measures. Since results were more pronounced in heat pain in the previous and current study, our analyses in the main manuscript focused on heat pain, but results for pressure pain are reported in the supplements.

### Evidence for fitness level, but not sex, effects on exercise-induced hypoalgesia

To identify whether exercise intensity affects pain ratings differently in the samples, we analysed the interaction of exercise intensity and subgroup (males from the previous study, females from the previous study, and females from the current study). We could show that females from the current study showed a significant difference between HI and LI exercise heat pain ratings at the highest pain intensity, but this was not evident in the males or females from the previous sample. To further evaluate the moderating roles of fitness level and sex in exercise-induced pain modulation, we examined these factors across the pooled sample. Both frequentist and Bayesian analyses converged in suggesting that fitness level is associated with variability in exercise-induced pain modulation, whereas evidence for sex-dependent effects was limited. Specifically, individuals with higher fitness levels tended to show greater hypoalgesia following HI compared with LI exercise, while sex did not meaningfully contribute to explaining differences in exercise-induced pain modulation. These findings extend our previous exploratory observations and suggest that fitness level may be a more important determinant of exercise-induced pain modulation than sex. Given the remaining uncertainty surrounding the magnitude of the fitness-related effects, some uncertainty regarding the magnitude of this effect remains.

A multitude of findings on exercise-induced hypoalgesia have been derived from earlier studies with male-dominated samples with higher fitness levels (Geisler et al., 2019; Haier et al., 1981a; Janal et al., 1984; Scheef et al., 2012), and results on sex dependent effects in exercise-induced hypoalgesia are still equivocal (Rice et al., 2019). Generally, (repeated) aerobic exercise has been shown to be associated with the activation of endogenous neuromodulatory systems (Gajic et al., 2026) such as endogenous opioid (Koltyn et al., 2014; Saanijoki et al., 2022), endocannabinoid (Matei et al., 2023), serotonergic and noradrenergic pathways (Anderson and Shivakumar, 2013; Beserra et al., 2018). Concerning the serotonergic and noradrenergic pathways, repeated aerobic exercise induces regulatory adaptations in glucocorticoid receptors in the hippocampus, prefrontal cortex and thalamus that potentially contribute to more efficient hypothalamic-pituitary-adrenal (HPA) axis regulation and reduced cortisol reactivity (Beserra et al., 2018). Likewise, positron emission tomography (PET) studies suggest that a higher aerobic fitness level is associated with reduced MOR binding in cortical (ACC, insula) and subcortical (ventral striatum) areas that are part of the descending pain modulatory system (DPMS) as well, suggesting that trained individuals show greater endogenous opioid release after aerobic exercise (Saanijoki et al., 2022). Overall, repeated aerobic exercise, resulting in a higher fitness level, might contribute to shaping (i.e. by increasing receptor density) or more strongly engaging already existing endogenous neuromodulatory systems and, thus, altering pain perception (Gajic et al., 2026; Song et al., 2022).

Concerning the mechanisms of sex potentially influencing exercise-induced pain modulation, the literature remains scarce and rather inconclusive. Since exercise has been linked to the release of endogenous opioids (Haier et al., 1981b; Janal et al., 1984) and, more recently, endocannabinoids (Crombie et al., 2018; Siebers et al., 2022, 2021), sex differences in these neuromodulatory systems might also contribute to the different expressions of exercise-induced pain modulation between the sexes. In terms of endogenous opioid signalling, previous studies indicate that men show larger mu-opioid receptor (MOR) activation in pain-related regions than women during sustained pain (Zubieta et al., 2002). Furthermore, sex differences in opioid analgesia remain controversial (Fullerton et al., 2018), pointing towards sex differences in the descending antinociceptive circuit. Specifically in the PAG, male rats showed higher MOR expression and binding compared with female rats (Loyd et al., 2008). Interestingly, increasing evidence in rodents also points towards sex differences in the expression and function of the endocannabinoid system in pain (Blanton et al., 2021), such as lower CB1 receptor density in the prefrontal cortex and amygdala (Castelli et al., 2014) or lower 2-AG concentration in the PAG (Levine et al., 2021) of female than male rodents. These regions are critical parts of the DPMS (Bingel and Tracey, 2008; Borras et al., 2004; Eippert et al., 2009; Petrovic et al., 2002) and are also involved in mediating exercise-induced pain modulation (Geisler et al., 2019; Nold et al., 2025; Scheef et al., 2012).

Of those studies that have included both sexes in their samples (Naugle et al., 2014; Niwa et al., 2022), a dose-response relationship of aerobic exercise intensity and the analgesic response could be shown in the overall sample, where high-intensity (70% heart rate reserve (HRR)) aerobic exercise produced larger hypoalgesic effects compared to moderate-intensity (50% HRR) exercise (Naugle et al., 2014; Niwa et al., 2022) and low-intensity (30% HRR) exercise (Niwa et al., 2022). Importantly, despite including both sexes, these studies did not include sex as a factor in their analyses, potentially confounding the observed effects (Tesarz et al., 2012). Of those studies that have explicitly accounted for sex as a moderating variable in exercise-induced hypoalgesia, some studies could show a hypoalgesic response from exercise in both sexes (Koltyn et al., 2014, 2001), whereas others (Gajsar et al., 2017; Lemley et al., 2016) found a stronger hypoalgesic response in females. When administering an aerobic exercise protocol, one study found the hypoalgesic response following exercise to be more pronounced in females compared to males (Vaegter et al., 2014).

Despite some studies showing comparable or greater hypoalgesic effects in females, these studies rarely account for fitness levels or athletic status (Tesarz et al., 2012). Generally, it has been shown that athletes exhibit increased pain tolerance and decreased pain intensity compared to non-athletic controls (Geva and Defrin, 2013; Tesarz et al., 2012). Among the few studies that have explicitly considered both fitness levels and sex in the hypoalgesic response to exercise, one study (Sternberg et al., 2001) found that treadmill running induced hypoalgesia in the cold pressor test in females, regardless of athletic status, but not in males. Notably, both male and female athletes showed a significant decrease in forearm withdrawal latency from a radiant heat device after treadmill exercise (Sternberg et al., 2001). Further corroborating this, female athletes experienced more pronounced hypoalgesia following aerobic exercise, especially during cold pressor and thermal pain tests (Smith, 2004). For isometric exercise, one study (Black et al., 2017) found that females demonstrated analgesic responses regardless of their physical activity levels, which is in contrast to our previous findings (Nold et al., 2025), where females did not show an analgesic response after aerobic exercise. Thus, the extent of sex differences might also vary depending on the type of exercise, the pain stimulus administered, and the measurement method employed (Tesarz et al., 2012), emphasising the complexity of sex-specific pain modulation mechanisms. Overall, female athletes tend to report pain similarly to their male counterparts (Tesarz et al., 2012), suggesting that athletic training may mitigate typical sex differences in the hypoalgesic response after exercise and potentially reduce disparities observed in the general population. Our results advance this by showing that fitness level, but not sex, moderates the hypoalgesic response to heat pain, even after short bouts of aerobic exercise.

## Limitations

On the experimental day, heat stimuli with a target intensity of VAS 30 were rated below the pain threshold. The anticipated intensities of VAS 50 and VAS 70 were rated above the pain threshold, suggesting that they were perceived as painful. However, the lower pain ratings on the experimental day as opposed to the calibration day are likely due to habituation effects between days (Ellerbrock et al., 2015; May et al., 2012). Furthermore, despite assessing exercise-related expectations and identifying participants to expect decreased whole-body pain following exercise, we did not experimentally manipulate those expectations, precluding causal conclusions regarding their relative contribution to the observed effects. Although expectations can modulate pain perception (Colloca et al., 2018; Jones et al., 2017), positive exercise-related expectations alone do not appear to augment exercise-induced hypoalgesia (Vaegter et al., 2020), suggesting that exercise-related expectations do not account for our findings. Moreover, as we compared HI and LI exercise but no sedentary rest condition, we cannot draw conclusions on the overall effect of exercise as opposed to no exercise on pain. Thus, it could be possible that even LI exercise suffices to evoke a hypoalgesic response (Niwa et al., 2022) or that individuals with higher fitness levels can show a distinction in the analgesic response based on the exercise intensity. However, previous research has suggested that HI exercise produces greater hypoalgesia compared to LI exercise (Jones et al., 2019; Naugle et al., 2014; Niwa et al., 2022; Smith, 2004; Vaegter et al., 2014, p. 201). Therefore, we chose the LI exercise as the control condition instead of rest to contribute to the understanding of the dose-response relationship between exercise intensities and hypoalgesic effects. Future research should investigate whether LI exercise suffices to induce an analgesic response.

## Conclusion

In this study, we could show that participants with relatively higher fitness levels (as measured by weight-corrected FTP) experience greater pain relief following high-intensity compared to low-intensity exercise, especially at high pain intensities. Together with our previous findings, these results suggest that fitness level may play a more central role than sex in explaining variability in exercise-induced pain modulation.

## Methods

### Participants

Overall, *N* = 22 female participants were recruited for this study from local cycling clubs and by (online) advertisements. Participants were required to be aged between 18 and 50 years and have a body mass index (BMI) ranging from 18 to 30, and to regularly participate in structured cycling training sessions per week. One participant was excluded due to a technical failure on the experimental day, resulting in incomplete data. For one participant, heat pain calibration failed, resulting in *N* = 20 participants for heat pain data and *N* = 21 participants for pressure pain data (for participant characteristics, see Table 1). It is crucial to note that the *N* = 21 females included in the current study did not participate in the previous study (Nold et al., 2025).

**Table 1.** Participant characteristics from the previous (Nold et al., 2025) and current study.

|  | Previous Study (Nold et al., 2025) |  |  | Current Study |
| --- | --- | --- | --- | --- |
|  | Overall<br>(*M*±*SD*) | Females<br>(*M*±*SD*) | Males<br>(*M*±*SD*) | Females<br>(*M*±*SD*) |
| *N* | 39 | 21 | 18 | 21 |
| Age (years) | 26.03 (4.83) | 25.33 (5.10) | 26.83 (4.35) | 32.29 (7.64) |
| Weight (kg) | 70.95 (12.14) | 63.33 (7.53) | 79.83 (10.38) | 63.6 (8.09) |
| Height (cm) | 177.10 (9.08) | 170.52 (6.35) | 184.78 (4.58) | 170 (5.95) |
| BMI | 22.51 (2.64) | 21.80 (2.59) | 23.33 (2.53) | 21 (1.93) |
| FTP (Watts/kg) | 1.79 (0.48) | 1.58 (0.44) | 2.03 (0.40) | 2.25 (0.41) |
| Training Volume (h/w) | 4.44 (3.24) | 3.88 (2.63) | 5.08 (3.81) | 13.10 (12.28) |

*M* = Mean; *SD* = Standard deviation; kg = kilogram; cm = centimetre; FTP = Functional Threshold Power. h/w = hours per week.

### Experimental Paradigm

The experimental paradigm was similar to that of the original study (Nold et al., 2025), using the same study setup and equipment (CPAR and TSA 2 and Wahoo KICKR bike). More precisely, participants completed four cycling blocks in total, two HI and two LI exercise blocks, presented in a pseudo-randomised order. Furthermore, the same calibration procedures were used to calibrate heat and pressure pain as well as the functional threshold power using the FTP_20_ test (Allen and Coggan, 2012, 2006) as implemented and validated in previous studies (Borszcz et al., 2018; McGrath et al., 2019; Nold et al., 2025). However, participants in the current study did not receive a pharmacological intervention and did not undergo fMRI scans; therefore, they received the painful stimuli outside the MR scanner whilst lying in a supine position. To monitor the painful stimuli more closely, participants provided online (continuous) ratings whilst the pain stimulus was ongoing on a VAS scale ranging from “no sensation” (0) to “almost unbearably painful” (150), with VAS 50 marking “minimally painful” (pain threshold). Furthermore, we administered the same questionnaires as described in the previous study. For a detailed description of the study procedures and experimental paradigm, please refer to Nold et al. (2025). Overall, the current study consisted of one calibration day (Day 1) and one experimental day (Day 2).

### Behavioural Data Acquisition

During cycling, heart rate (HR), maintained power (in watts), and cadence (in revolutions per minute (RPM)) were acquired continuously. In *N* = 7 participants, HR data were not recorded due to a technical failure, but were monitored throughout the cycling. The rating of perceived exertion (RPE) on the BORG scale (Borg, 1998), ranging from “no exertion” (6) to “maximal exertion” (20), was provided after each cycling block. Furthermore, the elapsed time between each cycling and pain within each block was five minutes to correspond with the mean time required in the previous study. Following the cycling, participants received a total of 18 pain stimuli, with 9 heat and 9 pressure stimuli applied in an alternating fashion and with their respective intensities (30, 50, 70 VAS) applied in a randomised order. Whilst the pain was ongoing, participants were asked to rate their currently perceived pain/non-painful sensation using the left and right buttons of a button box (Logitech). When their perception did not change, the cursor should remain at the same position. The online rating time exceeded the stimulus length by 2 seconds to capture changes in perception when the stimulus ramped down. The online ratings were sampled at 70 – 75 samples per second and interpolated at 0.9 seconds.

### Statistical Analyses

The behavioural statistical analyses were performed in MATLAB and RStudio (Version 2021.09.1). We used the *lmer* and *emmeans* functions from the *lme4* package (Version 1.1-35.1) to conduct linear mixed effect (LMER) (Bates et al., 2014; Hox et al., 2010) and post-hoc *t*-tests in R, respectively. Furthermore, the *rstatix* package (Kassambara, 2019) was used to conduct one and two-sample *t*-tests. For the Bayesian approach, all analyses were performed in R using the package *brms* and *rstan* that implements Bayesian multilevel regression models via Hamilton Monte Carlo sampling in Stan (Stan Development Team, 2024).

To investigate whether the distribution of menstrual cycle phases as well as hormonal contraceptives differed within and between days, χ2-tests were calculated. Furthermore, we evaluated whether there was a significant effect of expectation about acute aerobic exercise on different pain dimensions and calculated one-sample *t*-tests (two-tailed). Furthermore, we conducted paired samples *t*-tests comparing pre- and post-mood ratings on the dimensions of dejection, fatigue, discontent, and drive. To compare the FTP, self-reported fitness levels, training volume, age, weight, height, and BMI between the previous sample (males and females) and the current sample, we calculated two-sample *t*-tests. To evaluate the exercise parameters within the current study, watts and HR were averaged across time and blocks for each participant and for both exercise intensities. The RPE was also averaged across blocks for participants and exercise intensity. We calculated one-sample *t*-tests comparing the respective measures between HI and LI exercise.

### Linear Models

For visualisation purposes, we displayed the online ratings (averaged across subjects, blocks, trials, and stimulus intensities) and the SEM following HI and LI exercise for heat and pressure pain separately (Supplemental Fig. S7). For a more comprehensive understanding, the averaged online ratings at each stimulus intensity were visualised in the Supplemental Fig. S8. To statistically evaluate the online pain ratings and improve consistency with the post-stimulus pain ratings reported in the previous study (Nold et al., 2025), we calculated the maximum rating by averaging the online rating across the whole stimulus duration (0 – 17 seconds) and extracting the peak rating. The peak pain ratings across the whole stimulus duration would most closely correspond to the post-stimulus pain ratings provided in the previous study, where participants were asked to rate the most painful sensation throughout the stimulus duration. We calculated separate LMER models with the dependent variables overall pain ratings, heat pain ratings, and pressure pain ratings, respectively. All LMER models included trial and block as fixed effects and subject as a random effect. In a first LMER model, we included exercise intensity as a fixed effect; in a second LMER model, the stimulus intensity and exercise intensity, as well as their interaction, served as fixed effects. Finally, we conducted post-hoc *t*-tests and applied a Tukey adjustment to correct for multiple comparisons.

To gain a better understanding of the overall effect of exercise intensity, fitness level and sex across the current and previous study (Nold et al., 2025), we pooled the datasets. From the previous study, only data from the control (saline) condition were included in the current analyses. To allow for comparability of the previous and current study, as well as the subgroups (males of the previous study, females of the previous study and females of the current study), we rescaled the maximum pain ratings of the current study to a common scale of 0 – 100 to correspond with the previous study. To investigate the effect of exercise intensity and subgroup on pain ratings across all stimulus intensities, as well as at VAS 70 only, we conducted an LMER model. The interaction of exercise intensity and subgroup was used to assess whether the magnitude of exercise-induced pain modulation differed between the subgroups. To investigate the effect of fitness level, sex, and subgroup on exercise-induced pain modulation, we calculated participant-level difference scores (pain ratings following low-intensity exercise minus pain following high-intensity exercise), such that larger values reflected greater hypoalgesia following high-intensity exercise. Fitness level, fitness level × sex, and fitness level × subgroup were, respectively, included in three different linear models as predictors. The treatment order was included as a fixed effect in each model. Furthermore, we calculated the correlation coefficient using Pearson’s product-moment correlation for each of the respective models. These analyses correspond to the analyses conducted in Nold et al. (2025).

### Bayesian analyses

We conducted Bayesian analyses to complement and extend frequentist analyses by quantifying evidence for competing hierarchical models and estimating posterior distributions of regression coefficients (*β*) in the pooled samples model. All analyses modelled pain ratings as the dependent variable using Gaussian linear mixed-effects models. To account for repeated measurements within participants, all models included subject–specific intercepts. Furthermore, trial and block were included as fixed effects in all models. Models were organised in a hierarchical comparison framework consisting of a null model (including only nuisance covariates), main effects models (exercise intensity/VAS intensity) and interaction models (interaction terms). Overall, weakly informative priors were specified for all models (Table 2). Prior specifications were identical across models, with the only difference in intercepts depending on the ratings scale used (current study: 0-150; pooled dataset: 0-100) (for details, see Table 2). All models were estimated using four Markov chain Monte Carlo (MCMC) chains with 4000 iterations per chain (2000 warm-up). For more complex interaction models, adapt_delta was increased (up to 0.995), and maximum tree depth was increased (up to 15) to ensure stable sampling. Convergence was assessed using the potential scale reduction statistic (Ȓ ≈ 1.00), trace plots, and effective sample sizes.

**Table 2.**
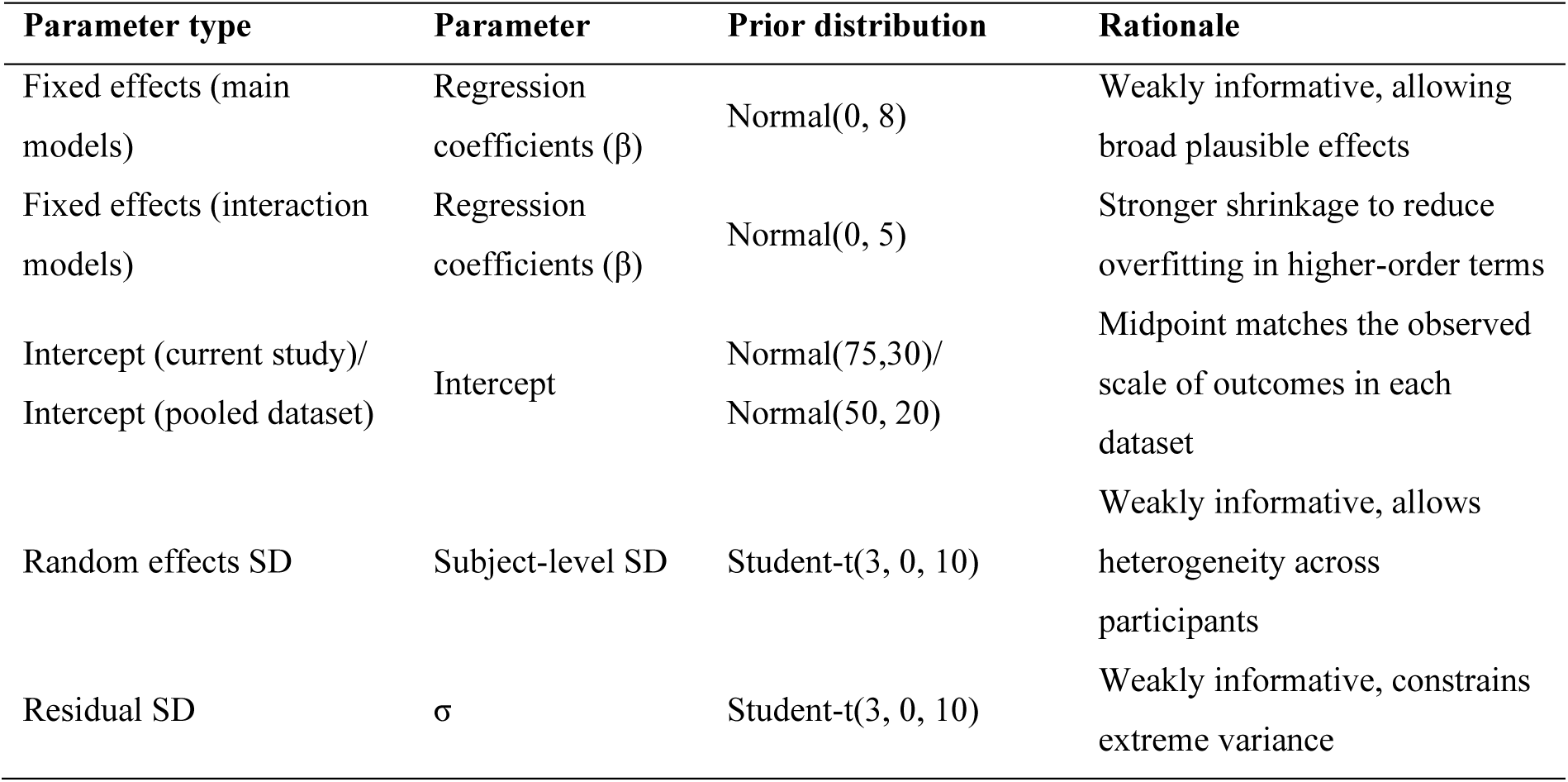
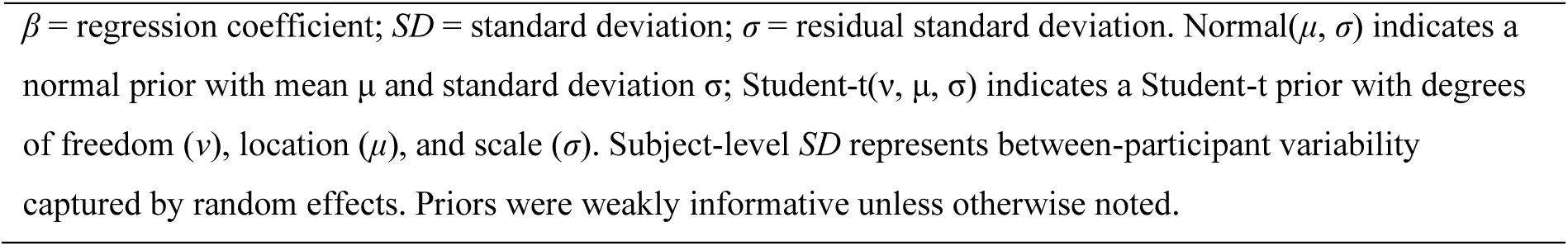
Prior specification for Bayesian mixed-effects models.

| Parameter type | Parameter | Prior distribution | Rationale |
| --- | --- | --- | --- |
| Fixed effects (main models) | Regression coefficients ($\beta$) | Normal(0, 8) | Weakly informative, allowing broad plausible effects |
| Fixed effects (interaction models) | Regression coefficients ($\beta$) | Normal(0, 5) | Stronger shrinkage to reduce overfitting in higher-order terms |
| Intercept (current study)/<br>Intercept (pooled dataset) | Intercept | Normal(75,30)/<br>Normal(50, 20) | Midpoint matches the observed scale of outcomes in each dataset |
| Random effects SD | Subject-level SD | Student-t(3, 0, 10) | Weakly informative, allows heterogeneity across participants |
| Residual SD | $\sigma$ | Student-t(3, 0, 10) | Weakly informative, constrains extreme variance |

$\beta$ = regression coefficient; $SD$ = standard deviation; $\sigma$ = residual standard deviation. $\text{Normal}(\mu, \sigma)$ indicates a normal prior with mean $\mu$ and standard deviation $\sigma$ ; $\text{Student-t}(\nu, \mu, \sigma)$ indicates a Student-t prior with degrees of freedom ( $\nu$ ), location ( $\mu$ ), and scale ( $\sigma$ ). Subject-level $SD$ represents between-participant variability captured by random effects. Priors were weakly informative unless otherwise noted.

### Bayesian model comparison

Bayesian model comparison was conducted using the Bayes Factor (BF_10_) estimated via bridge sampling for all frequentist linear model analyses. The Bayes Factor quantifies the relative evidence provided by the data for one model over another by comparing their marginal likelihoods. Specifically, BF_10_ expresses the evidence for the more complex model relative to a simpler model. Generally, BF_10_ > 1 indicates support for the more complex model over the simpler model and vice versa for BF_10_ < 1. According to the conventional evidence thresholds (Lovric, 2025), a 1 < BF_10_ < 3 indicates anecdotal/weak evidence, 3 < BF_10_ < 10 moderate evidence, and BF_10_ > 10 strong evidence in favour of the complex model. BF_10_ close to 1 indicates insensitive or inconclusive evidence for the complex over the simpler model. We computed the Bayes factor for the main effects model vs the null model, the interaction model vs. main effect model, and the full model vs. null model.

### Posterior inference

For the final analysis investigating the effect of exercise intensity, fitness level and sex in the pooled dataset, posterior inference was conducted. The model included pain ratings as the dependent variable with exercise intensity, fitness level, and sex, as well as subgroup as fixed effects, along with their interactions with exercise intensity. Again, trial and block were included as fixed effects along with subject-specific intercepts. Importantly, this model did not rely on the difference in pain ratings as in the frequentist analyses but on raw pain ratings. Therefore, this should be treated not as confirmatory but as complementary analyses. All posterior summaries were based on four MCMC chains with satisfactory convergence (Ȓ ≈ 1.00) and sufficient effective sample sizes across parameters. Unlike frequentist statistics, Bayesian inference does not rely on point estimates and p-values but rather a full posterior distribution of each model parameter, allowing for direct probability statements about effect size and direction of effects. Posterior summaries focused on the fixed-effect regression coefficients (*β*), specifically: (i) the main effect of exercise intensity, (ii) its moderation by fitness level, (iii) sex, and (iv) subgroup (males from the previous study, females from the previous study, females from the current study). Posterior samples were extracted and reported from the fitted model and used to compute and visualise in three complementary ways. First, the posterior point estimates were computed as the posterior mean of each fixed-effect regression coefficient (*β*) representing the expected regression coefficient under the posterior distribution. Second, uncertainty was summarised using 95% credible Intervals, defined as 2.5% and 97.5% quantiles of the posterior distribution. These intervals indicate the range containing 95% of the posterior probability for the regression coefficient. Third, the posterior probabilities of direction were computed for the fixed effect regression coefficients (*β*) by calculating the proportion of posterior samples below (*Pr(β < 0)*) for the main effect of exercise intensity, and its interaction terms with fitness level, sex and subgroup. The posterior probabilities provide a direct probability statement about the direction of each effect.

## Acknowledgments

We thank the participants for taking part in the study. Further, we thank Alexandra Tinnermann for her helpful comments.

## Author Contributions

Conceptualisation, C.B. and J.N.; Methodology, C.B. and J.N.; Investigation, C.B., J.N., T.F., and Z.G.; Visualisation, J.N.; Writing – Original Draft, C.B., J.N., and Z.G.; Funding Acquisition, C.B.; Resources, C.B.; Supervision, C.B.

## Ethics

Human subjects: All participants gave informed written consent. The study was approved by the Ethics Board of the Hamburg Medical Association (PV7456/2020-10144-BO-ff). We support inclusive, diverse, and equitable conduct of research.

## Additional Files

Supplemental Information: Supplemental Figures S1–S8 and Supplemental Tables S1-S30.

## Data availability

The dataset for the raw behavioural data generated in the current study is available from the corresponding author on reasonable request and after publication. All necessary data to evaluate the results of the study are included in the manuscript and supplementary materials.

## Code availability

This study was programmed using MATLAB 2021b and Psychophysics Toolbox (Version 3.0.19). For data visualisation, we used RStudio (Version 2026.05.0). The custom behavioural analysis pipelines are available on the public repository https://github.com/jannenold/hotspin_analyses_behavioural.git after publication.

