## Supplemental Materials for "Fitness Level, but not Sex, affects Exercise-Induced Pain Modulation"

*Lead contact and corresponding author.

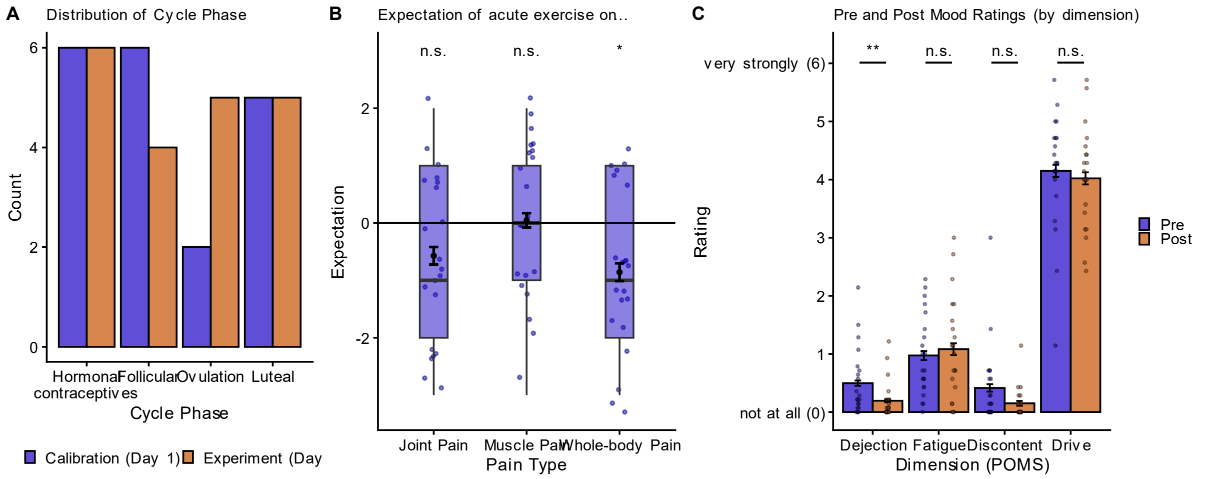


**Figure S1**. Evaluation of menstrual cycle phase, expectation, and mood. (**A**) Distribution of the cycle phase yielded no significant difference across ($\chi2(3) = 1.66$, $P = 0.65$) as well as within days (Day 1: $\chi2(3) = 2.26$, $P = 0.52$; Day 2: $\chi2(3) = 0.40$, $P = 0.94$), suggesting an equal distribution of cycle phases and intake of hormonal contraceptives in the sample. (**B**) Expectations of acute aerobic exercise on different pain types (joint pain, muscle pain, whole-body pain) ranged from 'greatly increase' (3) to 'greatly reduce' (-3) to capture the direction and magnitude of potential expectation effects. One sample *t*-test (two-tailed) revealed that participants did not expect acute aerobic exercise to change ($t(20) = -1.78$, $P = 0.09$) or muscle pain ($t(20) = 0.15$, $P = 0.88$), but to significantly reduce whole-body pain ($t(20) = -2.83$, $P = 0.01$). Dots depict subject-specific ratings, and solid black dots and error bars represent mean and SEM ($N = 21$). Boxplots show the distribution of data. (**C**) There were no significant differences in pre (blue) and post (orange) mood ratings (provided before and after the experimental day, respectively) in the dimensions fatigue ($t(20) = -0.50$, $P = 0.62$), discontent ($t(20) = 1.84$, $P = 0.08$), and drive ($t(20) = 0.58$, $P = 0.57$) but dejection was significantly decreased post compared to pre ($t(20) = 3.61$, $P = 0.002$). Dots depict subject-specific ratings. Barplot and error bars represent mean and SEM ($N = 21$).

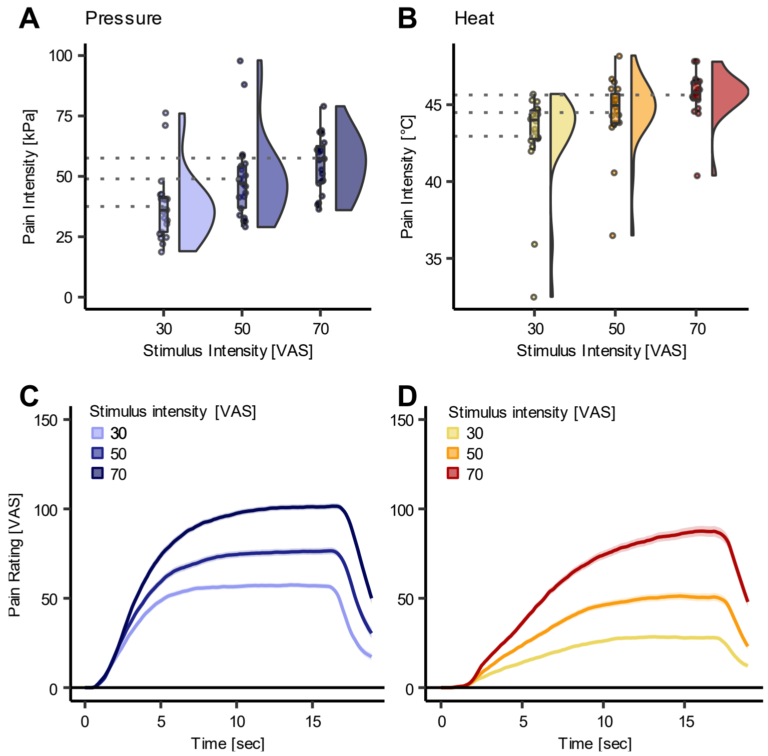


**Figure S2**. Raincloud plots for the distribution of the absolute intensities of the calibration of (**A**) pressure (kilopascal (kPa)) and (**B**) heat pain (in °C) at stimulus intensities corresponding to 30, 50, and 70 on a visual analogue scale (VAS). (**C, D**) Behavioural online ratings for (**C**) pressure and (**D**) heat pain throughout the stimulus duration (averaged across subjects and exercise intensities) at intensities corresponding to calibrated VAS 30, 50, and 70. For the online rating, the scale ranged from 'no sensation' to 'almost unbearably painful'. A rating of VAS 50 on the online rating scale would correspond to the pain threshold, and a rating of VAS 150 to 'almost unbearably painful'.

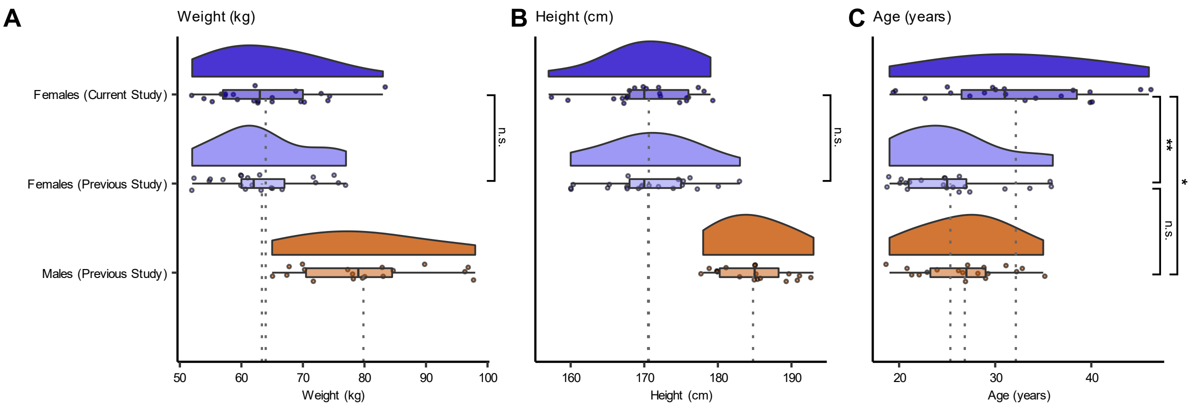


**Figure S3.** Comparing participant characteristics of the females of the current study (*N* = 21; dark blue) with the females (*N* = 21; light blue) and the males (*N* = 18; red) of the previous study (Nold et al., 2025). No significant difference in (**A**) weight *t*(40) = 0.26, *P* = 0.80) and (**B**) height (t(40) = 0.05, *P* = 0.96) between females in the current and previous study. Thus, BMI did not differ significantly between female participants across the two studies (*t*(36.87) = 0.15, *P* = 0.88). (**C**) Age differed significantly between females of both studies (t(32.98) = 3.27, *P* = 0.003) as well as the females of the current study and the males of the previous study (t(30.81) = 2.61, *P* = 0.01). Raincloud plots depict the distribution, and dots depict subject-specific data. n.s. = not significant, * *P* < 0.05, ** *P* < 0.01, *** *P* < 0.001.

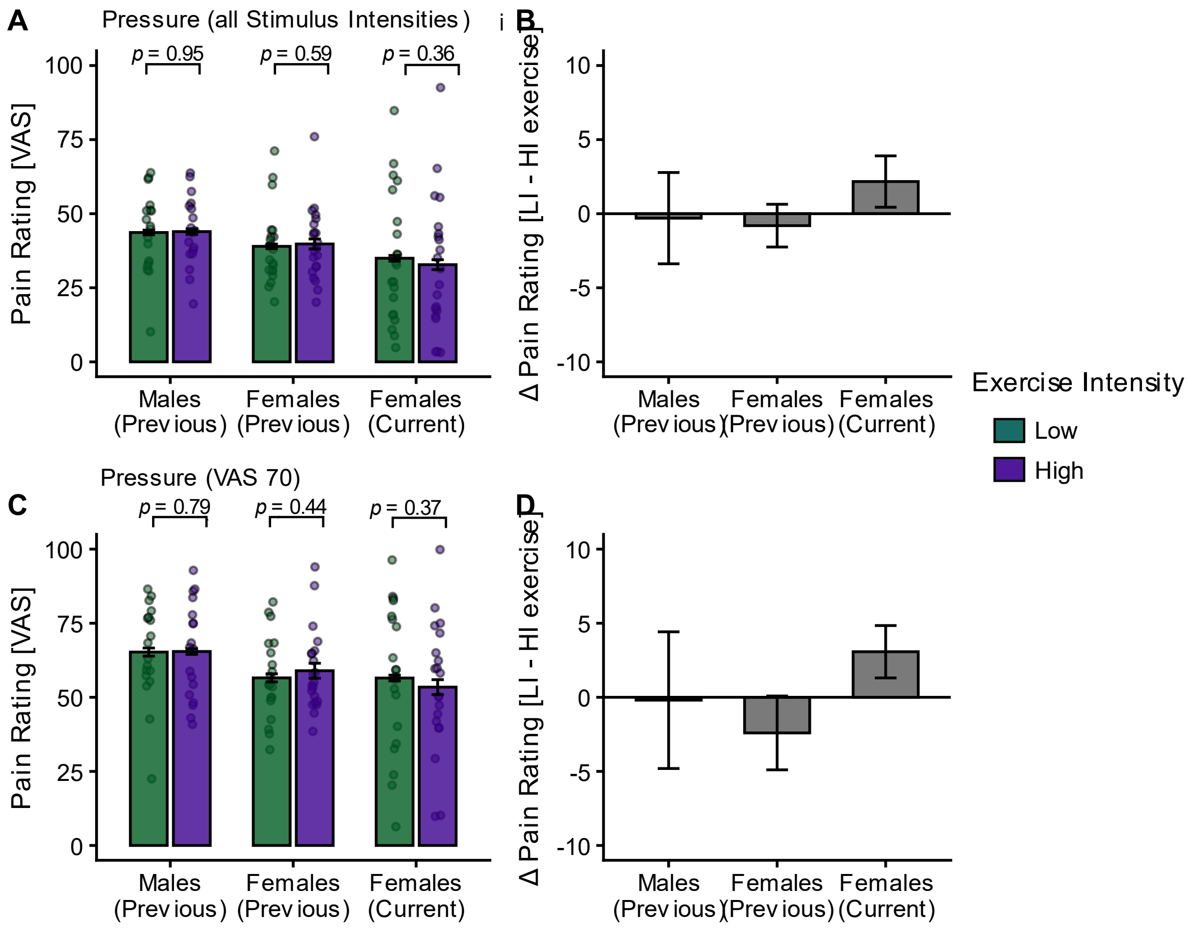


**Figure S4**. The effect of exercise intensity and subgroup (males from the previous study, females from the previous study and females from the current study) on pressure pain ratings. (**A**) Pressure pain ratings averaged across all stimulus intensities (VAS 30, 50, and 70) in each subgroup for LI (green) and HI (purple) exercise. There was no significant interaction of exercise intensity and subgroup on pressure pain ratings ($P$ = 0.52). (**B**) Differences between HI and LI exercise (LI – HI) of pain ratings across all stimulus intensities are visualised for each subgroup. (**C**) Pressure pain ratings at VAS 70 in each subgroup for LI (green) and HI (purple) exercise. There was no significant interaction of exercise intensity and subgroup on pressure pain ratings ($P$ = 0.44). (**D**) Differences between HI and LI exercise (LI – HI) of pain ratings at VAS 70 are visualised for each subgroup. Dots depict subject-specific pain ratings averaged across trials and blocks. P-values were calculated using post-hoc t-tests for the respective LMER models. Error bars depict the SEM (N = 60). The full model outputs can be found in Supplemental Tables S15 – S18.

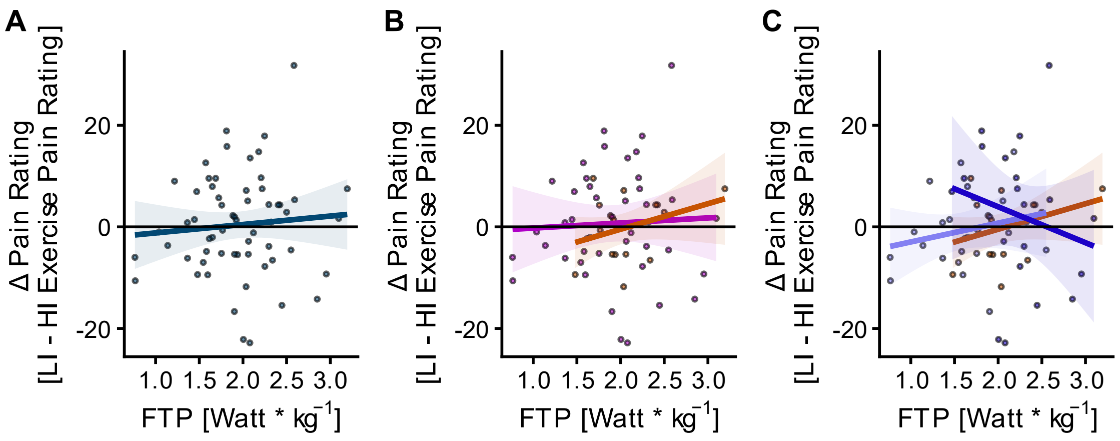


**Figure S5**. Effect of fitness level, sex, and subgroup on subject-specific differences in pressure pain ratings (dots) between low-intensity (LI) and high-intensity (HI) exercise conditions (LI – HI exercise pain ratings) pooled across samples and all stimulus intensities. (**A**) Fitness level (measured as functional threshold power; FTP) showed no significant main effect on difference pain ratings (*P* = 0.84) with a non-significant relation (*r* = 0.08, *P* = 0.52). (**B**) No significant interaction of fitness level × sex on the difference pain ratings (*P* = 0.48), with a non-significant relation in males (red line; *r* = 0.34, *P* = 0.17) and in females (pink line; pooled across current and previous studies; *r* = 0.05, *P* = 0.76). (**C**) No significant interaction fitness level × subgroup (*P* = 0.16) with non-significant correlations for females from the previous (*r* = 0.21, *P* = 0.35; Fig. 5C, light blue line) and current study (*r* = -0.21, *P* = 0.37; Fig. 5C, dark blue line). Full LMER model outputs can be found in Supplemental Tables S22 – S24.

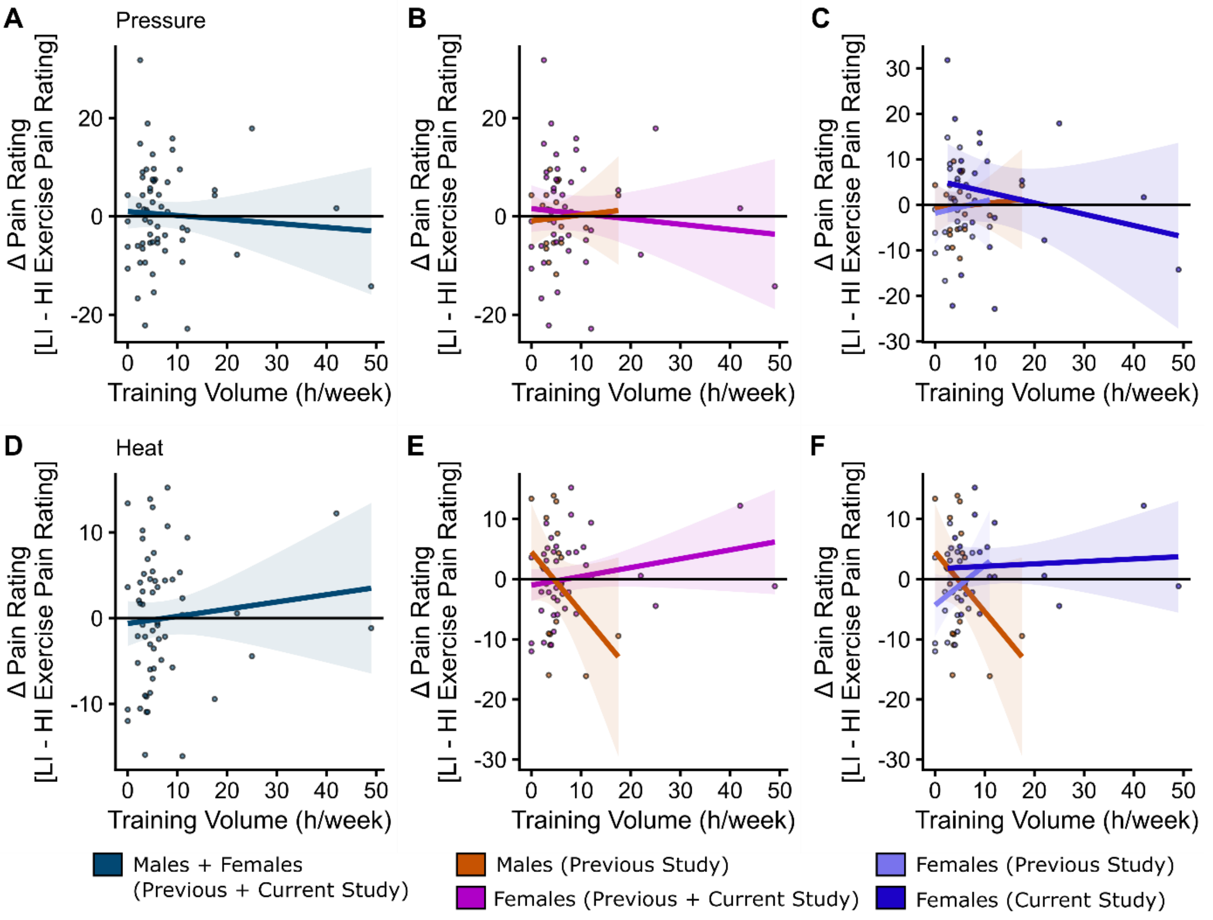


**Figure S6**. Effect of fitness level as measured as absolute training volume (hours/week), sex, and subgroup on subject-specific differences in pressure and heat pain ratings (dots) between low-intensity (LI) and high-intensity (HI) exercise conditions (LI – HI exercise pain ratings) pooled across samples and all stimulus intensities. (**A**) Fitness level (measured as training volume) showed no significant main effect on the difference pressure pain ratings (*P* = 0.36), with a non-significant relation (*r* = -0.07, *P* = 0.60). (**B**) No significant interaction of fitness level × sex on the difference pressure pain ratings (*P* = 0.73) with a non-significant relation in males (red line; *r* = 0.08, *P* = 0.77) and in females (pink line; pooled across current and previous study; *r* = -0.09, *P* = 0.56). (**C**) No significant interaction fitness level × subgroup (*P* = 0.52) with non-significant correlations for females from the previous (*r* = 0.09, *P* = 0.71; light blue line) and in current study (*r* = -0.21, *P* = 0.35; dark blue line). (**D**) Fitness level (measured as training volume) showed no significant main effect on difference heat pain ratings (*P* = 0.44) with a non-significant relation (*r* = 0.10, *P* = 0.47). (**E**) Significant interaction of fitness level × sex on difference heat pain ratings (*P* = 0.02) with a non-significant negative relation in males (red line; *r* = 0.12, *P* = -0.38) and a non-significant positive relation in females (pink line; pooled across current and previous study; *r* = -0.22, *P* = 0.16). (**F**) No significant interaction fitness level × subgroup (*P* = 0.13) with non-significant correlations for females from the previous (*r* = 0.27, *P* = 0.73; light blue line) and current study (*r* = -0.08, *P* = 0.73; dark blue line). Full LMER model outputs can be found in Supplementary Tables S25 – S30.

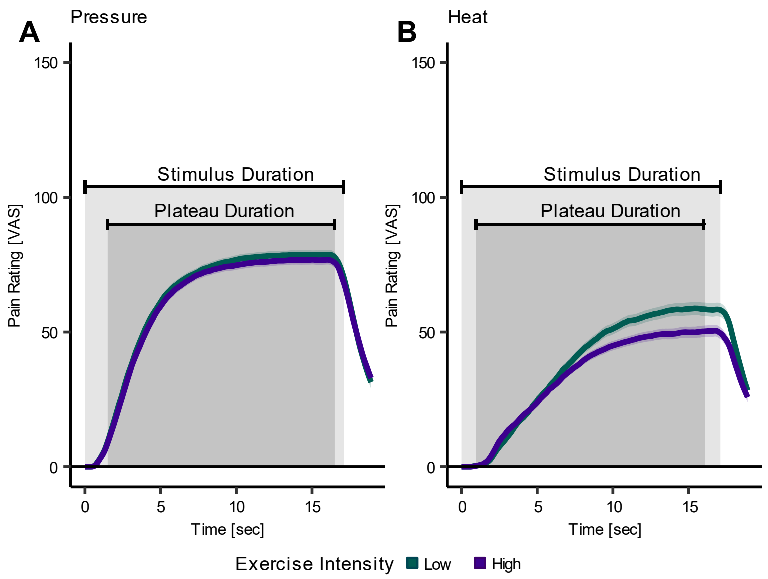


**Figure S7**. Behavioural online ratings for pressure and heat pain throughout the stimulus duration after high- (purple) and low- (green) intensity exercise (averaged across all stimulus intensities and participants). (**A**) No significant interaction of exercise intensity x time on pressure pain ratings ($\beta$ = -0.07, *SE* = 0.10, *t*(7956) = -0.71, *P* = 0.48). (**B**) Significant interaction of exercise intensity x time on heat pain ratings ($\beta$ = -0.13, *SE* = 0. 06, *t*(7577) = -2.12, *P* = 0.03). Online ratings were sampled at 70-75 samples/second throughout the stimulus duration (17 seconds) and 2 seconds after the stimulus end. The continuous ratings were interpolated at 0.9 seconds (1/2 of TR 1.8 seconds). Participants were instructed that the anchor points of the VAS scale correspond to "no perception" (0) and "almost unbearably painful" (150, i.e., pain tolerance). A rating of VAS 50 represents the pain threshold ("minimally painful"). The light and dark-shaded grey areas depict the stimulus duration (including ramp-up and ramp-down times) and the plateau duration (where constant temperature/kPa was applied), respectively. The shaded areas around the curves represent the SEM (pressure: *N* = 21, heat: *N* = 20).

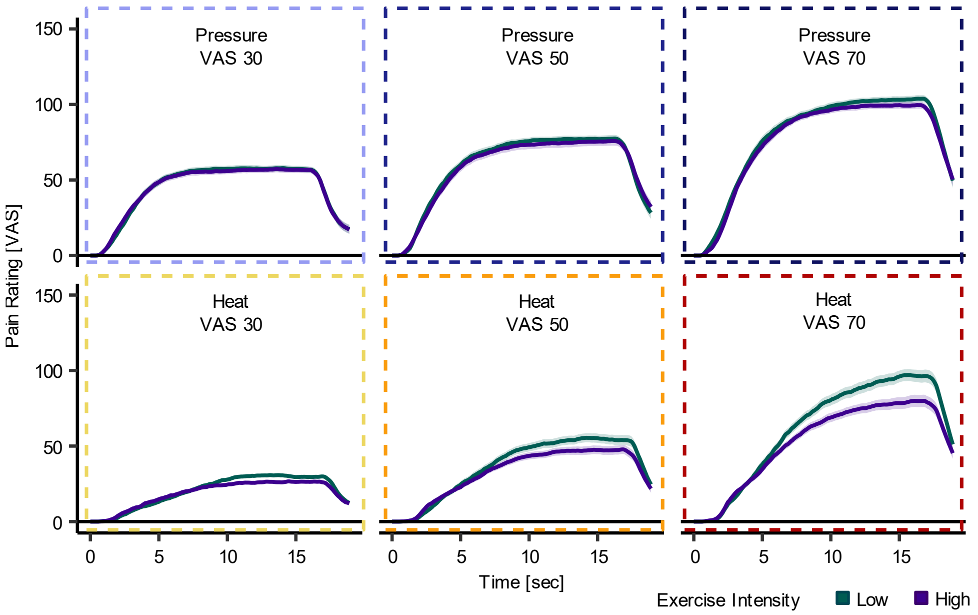


**Figure S8**. Behavioural online ratings for pressure (top) and heat (bottom) pain throughout the stimulus duration after high- (purple) and low- (green) intensity exercise at all stimulus intensities (VAS 30, 50, 70). No significant interaction of exercise intensity x time x stimulus intensity on pressure pain ratings ($\beta$ = -0.0002, *SE* = 0.004, *t*(2353) = -0.06, *P* = 0.95). Significant interaction of exercise intensity x time x stimulus intensity on heat pain ratings ($\beta$ = -0.01, *SE* = 0.003, *t*(2201) = -3.24, *P* = 0.001). Online ratings were sampled at 70-75 samples/second throughout the stimulus duration (17 seconds) and 2 seconds after the stimulus end. The continuous ratings were interpolated at 0.9 seconds (1/2 of TR 1.8 seconds). Participants were instructed that the anchor points of the VAS scale correspond to "no perception" (0) and "almost unbearably painful" (150, i.e., pain tolerance). A rating of VAS 50 represents the pain threshold ("minimally painful"). The light and dark-shaded grey areas depict the stimulus duration (including ramp-up and ramp-down times) and the plateau duration (where constant temperature/kPa was applied), respectively. The shaded areas around the curves represent the SEM (pressure: *N* = 21, heat: *N* = 20).

**Table S1**. Full LMER model output of exercise intensity and stimulus intensity on behavioural overall pain ratings.

| Fixed Effects | Estimate | *SE* | *df* | *t* | *P* |
|---|---|---|---|---|---|
| Intercept | 74.43 | 4.96 | 58.99 | 15.00 | $\mathbf{2\times10^{-16}}$ |
| Exercise Intensity | -3.29 | 2.14 | 1133.64 | -1.53 | 0.13 |
| Trial | -0.58 | 0.20 | 1121.78 | -2.94 | **0.003** |
| Block | 1.49 | 1.14 | 1130.36 | 1.30 | 0.19 |

LMER = linear mixed effects model, *SE* = standard error, *df* = degrees of freedom. Subject was included as random effect.  Trial and block were included as fixed effects. *N* = 21.

**Table S2**. Full LMER model output of exercise intensity and stimulus intensity on behavioural pressure pain ratings.

| Fixed Effects | Estimate | *SE* | *df* | *t* | *P* |
|---|---|---|---|---|---|
| Intercept | 85.25 | 6.36 | 37.20 | 13.40 | **$8.19 \times 10^{-13}$** |
| Exercise Intensity | -3.05 | 2.15 | 570.12 | -1.42 | 0.16 |
| Trial | -0.63 | 0.20 | 567.02 | -3.15 | **0.002** |
| Block | 1.59 | 1.14 | 569.48 | 1.39 | 0.16 |

LMER = linear mixed effects model, *SE* = standard error, *df* = degrees of freedom. Subject was included as random effect. Trial and block were included as fixed effects. *N* = 21.

**Table S3**. Post-hoc *t*-tests comparing exercise intensities (LI and HI) for pressure pain (across all stimulus intensities).

| Fixed Effects | T | *SE* | *df* | *t-ratio* | *P* |
|---|---|---|---|---|---|
| LI vs. HI | 3.05 | 2.15 | 570 | 1.42 | 0.16 |

*SE* = standard error, *df* = degrees of freedom. LI = low-intensity, HI = high-intensity.

**Table S4**. Full LMER model output of exercise intensity and stimulus intensity on behavioural heat pain ratings.

| Fixed Effects | Estimate | *SE* | *df* | *t* | *P* |
|---|---|---|---|---|---|
| Intercept | 64.23 | 6.67 | 54.04 | 9.64 | **$2.47\times10^{-11}$** |
| Exercise Intensity | -4.13 | 2.93 | 538.70 | -1.41 | 0.16 |
| Trial | -0.47 | 0.27 | 531.96 | -1.76 | 0.08 |
| Block | 1.01 | 1.56 | 535.90 | 0.65 | 0.52 |

LMER = linear mixed effects model, *SE* = standard error, *df* = degrees of freedom. Subject was included as random effect. Trial and block were included as fixed effects. *N* = 20.

**Table S5**. Post-hoc *t*-tests comparing exercise intensities (LI and HI) for heat pain (across all stimulus intensities).

| Fixed Effects | T | *SE* | *df* | *t-ratio* | *P* |
|---|---|---|---|---|---|
| LI vs. HI | ´4.13 | 2.93 | 539 | 1.41 | 0.16 |

*SE* = standard error, *df* = degrees of freedom. LI = low-intensity, HI = high-intensity.

**Table S6**. Full LMER model output of exercise intensity and stimulus intensity on behavioural overall pain ratings.

| Fixed Effects | Estimate | *SE* | *df* | *t* | *P* |
|---|---|---|---|---|---|
| Intercept | 4.70 | 5.84 | 100.88 | 0.80 | 0.42 |
| Exercise Intensity | 7.76 | 5.23 | 1120.67 | 1.48 | 0.14 |
| Stimulus Intensity | 1.45 | 0.07 | 1120.31 | 19.62 | $\mathbf{<2\times10^{-16}}$ |
| Trial | -0.79 | 0.16 | 1119.86 | -5.08 | $\mathbf{4.45\times10^{-7}}$ |
| Block | 1.55 | 0.90 | 1125.10 | 1.74 | 0.08 |
| Stimulus Intensity x Exercise Intensity | -0.22 | 0.10 | 1119.88 | -2.22 | **0.03** |

LMER = linear mixed effects model, *SE* = standard error, *df* = degrees of freedom. Subject was included as random effect. Trial and block were included as fixed effects. *N* = 21.

**Table S7**. Full LMER model output of exercise intensity and stimulus intensity on behavioural pressure pain ratings.

| Fixed Effects | Estimate | *SE* | *df* | *t* | *P* |
|---|---|---|---|---|---|
| Intercept | 31.70 | 6.61 | 48.23 | 4.79 | **$1.61 \times 10^{-5}$** |
| Exercise Intensity | 1.27 | 4.72 | 565.15 | 0.27 | 0.79 |
| Stimulus Intensity | 1.13 | 0.07 | 565.20 | 17.04 | **$<2 \times 10^{-16}$** |
| Trial | -0.91 | 0.14 | 565.02 | -6.44 | **$2.61 \times 10^{-10}$** |
| Block | 1.60 | 0.81 | 566.32 | 2.00 | **0.05** |
| Stimulus Intensity x Exercise Intensity | -0.09 | 0.09 | 565 | -0.97 | 0.33 |

LMER = linear mixed effects model, *SE* = standard error, *df* = degrees of freedom. Subject was included as random effect. Trial and block were included as fixed effects. *N* = 21.

**Table S8**. Post-hoc *t*-tests (Tukey-adj.) for LMER models in pressure pain comparing the averaged pain ratings at HI and LI exercise at each stimulus intensity.

| | *T* | *SE* | *df* | *T-ratio* | *P* |
|---|---|---|---|---|---|
| VAS 30 | 1.73 | 2.54 | 564 | 0.68 | 0.50 |
| VAS 50 | 2.38 | 2.54 | 564 | 0.94 | 0.35 |
| VAS 70 | 5.27 | 2.63 | 564 | 2.01 | **0.05** |

SE = standard error, df = degrees of freedom. LI = low-intensity, HI = high-intensity.

**Table S9.** Full LMER model output of exercise intensity and stimulus intensity on behavioural heat pain ratings.

| Fixed Effects | Estimate | *SE* | *df* | *t* | *P* |
|---|---|---|---|---|---|
| Intercept | -21.39 | 7.23 | 58.68 | -2.96 | **0.004** |
| Exercise Intensity | 14.03 | 5.71 | 530.24 | 2.46 | **0.01** |
| Stimulus Intensity | 1.76 | 0.08 | 530.58 | 21.64 | **$<2\times10^{-16}$** |
| Trial | -0.54 | 0.17 | 529.92 | -3.19 | **0.001** |
| Block | 1.08 | 0.99 | 531.34 | 1.10 | 0.27 |
| Stimulus Intensity x Exercise Intensity | -0.36 | 0.11 | 529.95 | -3.32 | **0.001** |

LMER = linear mixed effects model, *SE* = standard error, *df* = degrees of freedom. Subject was included as random effect. Trial and block were included as fixed effects. *N* = 20.

**Table S10**. Post-hoc *t*-tests (Tukey-adj.) for LMER models in heat pain comparing the pain ratings at HI and LI exercise at each stimulus intensity.

| | *T* | *SE* | *df* | *T-ratio* | *P* |
|---|---|---|---|---|---|
| VAS 30 | -2.78 | 3.00 | 529 | -0.93 | 0.35 |
| VAS 50 | 3.37 | 3.00 | 529 | 1.13 | 0.26 |
| VAS 70 | 12.02 | 3.19 | 529 | 3.77 | **0.0002** |

SE = standard error, df = degrees of freedom. LI = low-intensity, HI = high-intensity.

**Table S11.** Full LMER model output of exercise intensity and subgroup on behavioural heat pain ratings (averaged across all stimulus intensities).

| Fixed Effects | Estimate | *SE* | *df* | *t* | *P* |
|---|---|---|---|---|---|
| Intercept | 55.96 | 5.94 | 115.57 | 9.42 | **$5.59\times10^{-16}$** |
| Exercise intensity | 3.01 | 3.52 | 1886.00 | 0.86 | 0.392 |
| Group | -11.11 | 2.70 | 69.06 | -4.11 | **0.0001** |
| Treatment order | 5.57 | 4.21 | 54.00 | 1.32 | 0.191 |
| Exercise intensity × Group | -1.63 | 1.68 | 1897.74 | -0.97 | 0.332 |

LMER = linear mixed effects model, *SE* = standard error, *df* = degrees of freedom. Subject was included as random effect. Trial, block, and treatment order were included as fixed effects. *N* = 59. Subgroup was defined as factor with levels males from previous study, females from previous study, and females from current study.

**Table S12**. Post-hoc *t*-tests (Tukey-adj.) for LMER models in heat pain (averaged across all stimulus intensities) comparing the pain ratings at HI and LI exercise for each subgroup.

| | *T* | *SE* | *df* | *T-ratio* | *P* |
|---|---|---|---|---|---|
| Males (previous study) | -0.34 | 2.23 | 1875 | -0.15 | 0.88 |
| Females (previous study) | -1.62 | 2.05 | 1875 | -0.79 | 0.43 |
| Females (current study) | 3.31 | 2.49 | 1918 | 1.33 | 0.18 |

SE = standard error, df = degrees of freedom. LI = low-intensity, HI = high-intensity. Males (previous study): *n* = 18; females (previous study): *n* = 21; females (current study): *n* = 20.

**Table S13.** Full LMER model output of exercise intensity and subgroup on behavioural heat pain ratings (VAS 70).

| Fixed Effects | Estimate | *SE* | *df* | *t* | *P* |
|---|---|---|---|---|---|
| Intercept | 83.61 | 8.41 | 70.56 | 9.94 | **$4.77\times10^{-15}$** |
| Exercise intensity | 11.12 | 3.16 | 560.87 | 3.52 | **0.0005** |
| Subgroup | -14.76 | 4.17 | 57.84 | -3.54 | **0.0008** |
| Treatment order | 10.02 | 6.66 | 53.27 | 1.50 | 0.14 |
| Exercise intensity × Group | -5.96 | 1.53 | 562.24 | -3.89 | **0.0001** |

LMER = linear mixed effects model, *SE* = standard error, *df* = degrees of freedom. Subject was included as random effect. Trial, block, and treatment order were included as fixed effects. *N* = 59. Subgroup was defined as factor with levels males from previous study, females from previous study, and females from current study.

**Table S14**. Post-hoc *t*-tests (Tukey-adj.) for LMER models in heat pain (VAS 70) comparing the pain ratings at HI and LI exercise for each subgroup.

| | *T* | *SE* | *df* | *T-ratio* | *P* |
|---|---|---|---|---|---|
| Males (previous study) | -4.35 | 1.97 | 559 | -2.20 | **0.03** |
| Females (previous study) | -0.69 | 1.81 | 559 | 2.90 | 0.71 |
| Females (current study) | 7.95 | 2.32 | 564 | 3.42 | **0.0007** |

SE = standard error, df = degrees of freedom. LI = low-intensity, HI = high-intensity. Males (previous study): *n* = 18; females (previous study): *n* = 21; females (current study): *n* = 20.

**Table S15.** Full LMER model output of exercise intensity and subgroup on behavioural pressure pain ratings (averaged across all stimulus intensities).

| Fixed Effects | Estimate | *SE* | *df* | *t* | *P* |
|---|---|---|---|---|---|
| Intercept | 46.91 | 6.22 | 83.34 | 7.55 | **$5.06\times10^{-11}$** |
| Exercise intensity | 1.54 | 2.79 | 1915.36 | 0.55 | 0.58 |
| Group | -5.34 | 2.99 | 63.00 | -1.79 | 0.08 |
| Treatment order | 3.74 | 4.84 | 55.78 | 0.77 | 0.44 |
| Exercise intensity × Group | -0.86 | 1.32 | 1921.03 | -0.65 | 0.52 |

LMER = linear mixed effects model, *SE* = standard error, *df* = degrees of freedom. Subject was included as random effect. Trial, block, and treatment order were included as fixed effects. *N* = 60. Subgroup was defined as factor with levels males from previous study, females from previous study, and females from current study.

**Table S16**. Post-hoc *t*-tests (Tukey-adj.) for LMER models in pressure pain (averaged across all stimulus intensities) comparing the pain ratings at HI and LI exercise for each subgroup.

| | *T* | *SE* | *df* | *T-ratio* | *P* |
|---|---|---|---|---|---|
| Males (previous study) | -0.10 | 1.77 | 1910 | -0.06 | 0.96 |
| Females (previous study) | -0.87 | 1.63 | 1910 | -0.53 | 0.59 |
| Females (current study) | 1.78 | 1.93 | 1929 | 0.92 | 0.36 |

SE = standard error, df = degrees of freedom. LI = low-intensity, HI = high-intensity. Males (previous study): *n* = 18; females (previous study): *n* = 21; females (current study): *n* = 21.

**Table S17.** Full LMER model output of exercise intensity and subgroup on behavioural pressure pain ratings (VAS 70).

| Fixed Effects | Estimate | *SE* | *df* | *t* | *P* |
|---|---|---|---|---|---|
| Intercept | 68.21 | 6.73 | 91.50 | 10.14 | **$<2\times10^{-16}$** |
| Exercise intensity | 2.56 | 3.34 | 576.85 | 0.77 | 0.44 |
| Group | -5.77 | 3.18 | 64.33 | -1.82 | 0.07 |
| Treatment order | 5.31 | 5.07 | 54.94 | 1.05 | 0.30 |
| Exercise intensity × Group | -1.22 | 1.59 | 579.24 | -0.77 | 0.44 |

LMER = linear mixed effects model, *SE* = standard error, *df* = degrees of freedom. Subject was included as random effect. Trial, block, and treatment order were included as fixed effects. *N* = 60. Subgroup was defined as factor with levels males from previous study, females from previous study, and females from current study.

**Table S18**. Post-hoc *t*-tests (Tukey-adj.) for LMER models in pressure pain (VAS 70) comparing the pain ratings at HI and LI exercise for each subgroup.

| | *T* | *SE* | *df* | *T-ratio* | *P* |
|---|---|---|---|---|---|
| Males (previous study) | -0.57 | 2.12 | 574 | -0.27 | 0.79 |
| Females (previous study) | -1.51 | 1.95 | 574 | -0.78 | 0.44 |
| Females (current study) | 2.11 | 2.34 | 582 | 0.90 | 0.37 |

SE = standard error, df = degrees of freedom. LI = low-intensity, HI = high-intensity. Males (previous study): *n* = 18; females (previous study): *n* = 21; females (current study): *n* = 21.

**Table S19.** Full linear model output from the model including FTP on difference score heat pain ratings (LI – HI exercise) in the pooled samples.

| Fixed Effects | Estimate | *SE* | *t* | *P* |
|---|---|---|---|---|
| Intercept | -8.34 | 3.84 | -2.17 | **0.03** |
| FTP | 4.99 | 2.05 | 2.43 | **0.02** |
| Treatment_order | -2.15 | 2.13 | -1.01 | 0.32 |

FTP = functional threshold power (weight-corrected), *SE* = standard error, *df* = degrees of freedom.

**Table S20.** Full linear model output from the model including FTP and sex on difference score heat pain ratings (LI – HI exercise) in the pooled samples.

| Fixed Effects | Estimate | *SE* | *t* | *P* |
|---|---|---|---|---|
| Intercept | -3.99 | 4.09 | -0.98 | 0.33 |
| FTP | 3.05 | 2.27 | 1.35 | 0.18 |
| Sex | -24.36 | 9.57 | -2.55 | **0.01** |
| Treatment_order | -2.27 | 2.18 | -1.04 | 0.30 |
| FTP x Sex | 11.12 | 4.69 | 2.37 | **0.02** |

FTP = functional threshold power (weight-corrected), *SE* = standard error, *df* = degrees of freedom.

**Table S21.** Full linear model output from the model including FTP and subgroup on difference score heat pain ratings (LI – HI exercise) in the pooled samples.

| Fixed Effects | Estimate | *SE* | *t* | *P* |
|---|---|---|---|---|
| Intercept | -46.10 | 12.70 | -3.63 | **0.0006** |
| FTP | 21.65 | 6.08 | 3.56 | **0.0008** |
| Subgroup | 19.01 | 6.04 | 3.15 | **0.003** |
| Treatment_order | -4.00 | 2.22 | -1.80 | 0.08 |
| FTP x Subgroup | -7.99 | 2.76 | -2.89 | **0.005** |

FTP = functional threshold power (weight-corrected), *SE* = standard error, *df* = degrees of freedom. Subgroup was defined as factor with levels males from previous study, females from previous study, and females from current study.

**Table S22.** Full linear model output from the model including FTP on difference score pressure pain ratings (LI – HI exercise) in the pooled samples.

| Fixed Effects | Estimate | *SE* | *t* | *P* |
|---|---|---|---|---|
| Intercept | -2.68 | 5.21 | -0.51 | 0.61 |
| FTP | 0.57 | 2.78 | 0.21 | 0.84 |
| Treatment_order | 3.08 | 2.92 | 1.06 | 0.30 |

FTP = functional threshold power (weight-corrected), *SE* = standard error, *df* = degrees of freedom.

**Table S23.** Full linear model output from the model including FTP and sex on difference score pressure pain ratings (LI – HI exercise) in the pooled samples.

| Fixed Effects | Estimate | *SE* | *t* | *P* |
|---|---|---|---|---|
| Intercept | -0.87 | 5.84 | -0.15 | 0.88 |
| FTP | -0.41 | 3.23 | -0.13 | 0.90 |
| Sex | -9.91 | 13.78 | -0.72 | 0.48 |
| Treatment_order | 3.27 | 3.14 | 1.04 | 0.30 |
| FTP x Sex | 4.85 | 6.74 | 0.72 | 0.48 |

FTP = functional threshold power (weight-corrected), *SE* = standard error, *df* = degrees of freedom.

**Table S24.** Full linear model output from the model including FTP and subgroup on difference score pressure pain ratings (LI – HI exercise) in the pooled samples.

| Fixed Effects | Estimate | *SE* | *t* | *P* |
|---|---|---|---|---|
| Intercept | -28.71 | 18.61 | -1.54 | 0.13 |
| FTP | 12.71 | 8.90 | 1.43 | 0.16 |
| Subgroup | 12.83 | 8.12 | 1.46 | 0.15 |
| Treatment_order | 2.48 | 3.26 | 0.76 | 0.45 |
| FTP x Subgroup | -5.78 | 4.02 | -1.44 | 0.16 |

FTP = functional threshold power (weight-corrected), *SE* = standard error, *df* = degrees of freedom. Subgroup was defined as factor with levels males from previous study, females from previous study, and females from current study.

**Table S25.** Full linear model output from the model including Training Volume on difference score heat pain ratings (LI – HI exercise) in the pooled samples.

| Fixed Effects | Estimate | *SE* | *t* | *P* |
|---|---|---|---|---|
| Intercept | -0.24 | 1.73 | -0.14 | 0.89 |
| Training Volume | 0.10 | 0.12 | 0.78 | 0.44 |
| Treatment_order | -0.75 | 2.16 | -0.35 | 0.73 |

Training Volume measured as training hours per week. *SE* = standard error, *df* = degrees of freedom.

**Table S26.** Full linear model output from the model including FTP and sex on difference score heat pain ratings (LI – HI exercise) in the pooled samples.

| Fixed Effects | Estimate | *SE* | *t* | *P* |
|---|---|---|---|---|
| Intercept | -0.45 | 2.00 | -0.23 | 0.82 |
| Training Volume | 0.16 | 0.12 | 1.30 | 0.20 |
| Sex | 5.33 | 3.40 | 1.57 | 0.12 |
| Treatment_order | -0.92 | 2.15 | -0.42 | 0.67 |
| Training Volume x Sex | -1.14 | 0.49 | -2.33 | **0.02** |

Training Volume measured as training hours per week. *SE* = standard error, *df* = degrees of freedom.

**Table S27.** Full linear model output from the model including FTP and subgroup on difference score heat pain ratings (LI – HI exercise) in the pooled samples.

| Fixed Effects | Estimate | *SE* | *t* | *P* |
|---|---|---|---|---|
| Intercept | 2.69 | 4.48 | 0.60 | 0.55 |
| Training Volume | -0.99 | 0.69 | -1.44 | 0.16 |
| Subgroup | -0.43 | 2.05 | -0.21 | 0.83 |
| Treatment_order | -1.67 | 2.34 | -0.71 | 0.48 |
| Training Volume x Subgroup | 0.37 | 0.24 | 1.54 | 0.13 |

Training Volume measured as training hours per week. *SE* = standard error, *df* = degrees of freedom. Subgroup was defined as factor with levels males from previous study, females from previous study, and females from current study.

**Table S28.** Full linear model output from the model including FTP on difference score pressure pain ratings (LI – HI exercise) in the pooled samples.

| Fixed Effects | Estimate | *SE* | *t* | *P* |
|---|---|---|---|---|
| Intercept | -1.09 | 2.24 | -0.49 | 0.63 |
| Training Volume | -0.14 | 0.16 | -0.92 | 0.36 |
| Treatment_order | 4.04 | 2.81 | 1.44 | 0.16 |

Training Volume measured as training hours per week. *SE* = standard error, *df* = degrees of freedom.

**Table S29.** Full linear model output from the model including FTP and sex on difference score pressure pain ratings (LI – HI exercise) in the pooled samples.

| Fixed Effects | Estimate | *SE* | *t* | *P* |
|---|---|---|---|---|
| Intercept | -0.78 | 2.72 | -0.29 | 0.77 |
| Training Volume | -0.16 | 0.17 | -0.97 | 0.34 |
| Sex | -1.64 | 4.63 | -0.35 | 0.73 |
| Treatment_order | 3.96 | 2.92 | 1.35 | 0.18 |
| Training Volume x Sex | 0.23 | 0.67 | 0.35 | 0.73 |

Training Volume measured as training hours per week. *SE* = standard error, *df* = degrees of freedom.

**Table S30.** Full linear model output from the model including FTP and subgroup on difference score pressure pain ratings (LI – HI exercise) in the pooled samples.

| Fixed Effects | Estimate | *SE* | *t* | *P* |
|---|---|---|---|---|
| Intercept | -5.65 | 5.99 | -0.94 | 0.35 |
| Training Volume | 0.41 | 0.92 | 0.45 | 0.66 |
| Subgroup | 2.72 | 2.74 | 0.83 | 0.41 |
| Treatment_order | 3.21 | 3.13 | 1.03 | 0.31 |
| Training Volume x Subgroup | -0.21 | 0.32 | -0.64 | 0.52 |

Training Volume measured as training hours per week. *SE* = standard error, *df* = degrees of freedom.
Subgroup was defined as factor with levels males from previous study, females from previous study, and females from current study.
